# *In vivo* evolution of a *Klebsiella pneumoniae* capsule defect promotes complement-mediated opsono-phagocytosis and persistence during recurrent infection

**DOI:** 10.1101/2023.05.31.542722

**Authors:** William Bain, Brian Ahn, Hernán F. Peñaloza, Christi L. McElheny, Nathanial Tolman, Rick van der Geest, Shekina Gonzalez-Ferrer, Nathalie Chen, Xiaojing An, Ria Hosuru, Mohammadreza Tabary, Erin Papke, Naina Kohli, Nauman Farooq, William Bachman, Tolani F. Olonisakin, Zeyu Xiong, Marissa P. Griffith, Mara Sullivan, Jonathan Franks, Mustapha M. Mustapha, Alina Iovleva, Tomeka Suber, Robert Q. Shanks, Viviana P. Ferreira, Donna B. Stolz, Daria Van Tyne, Yohei Doi, Janet S. Lee

## Abstract

*Klebsiella pneumoniae* carbapenemase-producing *K. pneumoniae* (KPC-Kp) bloodstream infections rarely overwhelm the host but are associated with high mortality. The complement system is a key host defense against bloodstream infection. However, there are varying reports of serum resistance among KPC-Kp isolates. We assessed growth of 59 KPC-Kp clinical isolates in human serum and found increased resistance in 16/59 (27%). We identified five genetically-related bloodstream isolates with varying serum resistance profiles collected from a single patient during an extended hospitalization marked by recurrent KPC-Kp bloodstream infections. We noted a loss-of-function mutation in the capsule biosynthesis gene, *wcaJ,* that emerged during infection was associated with decreased polysaccharide capsule content, and resistance to complement-mediated killing. Surprisingly, disruption of *wcaJ* increased deposition of complement proteins on the microbial surface compared to the wild-type strain and led to increased complement-mediated opsono-phagocytosis in human whole blood. Disabling opsono-phagocytosis in the airspaces of mice impaired *in vivo* control of the *wcaJ* loss-of-function mutant in an acute lung infection model. These findings describe the rise of a capsular mutation that promotes KPC-Kp persistence within the host by enabling co-existence of increased bloodstream fitness and reduced tissue virulence.

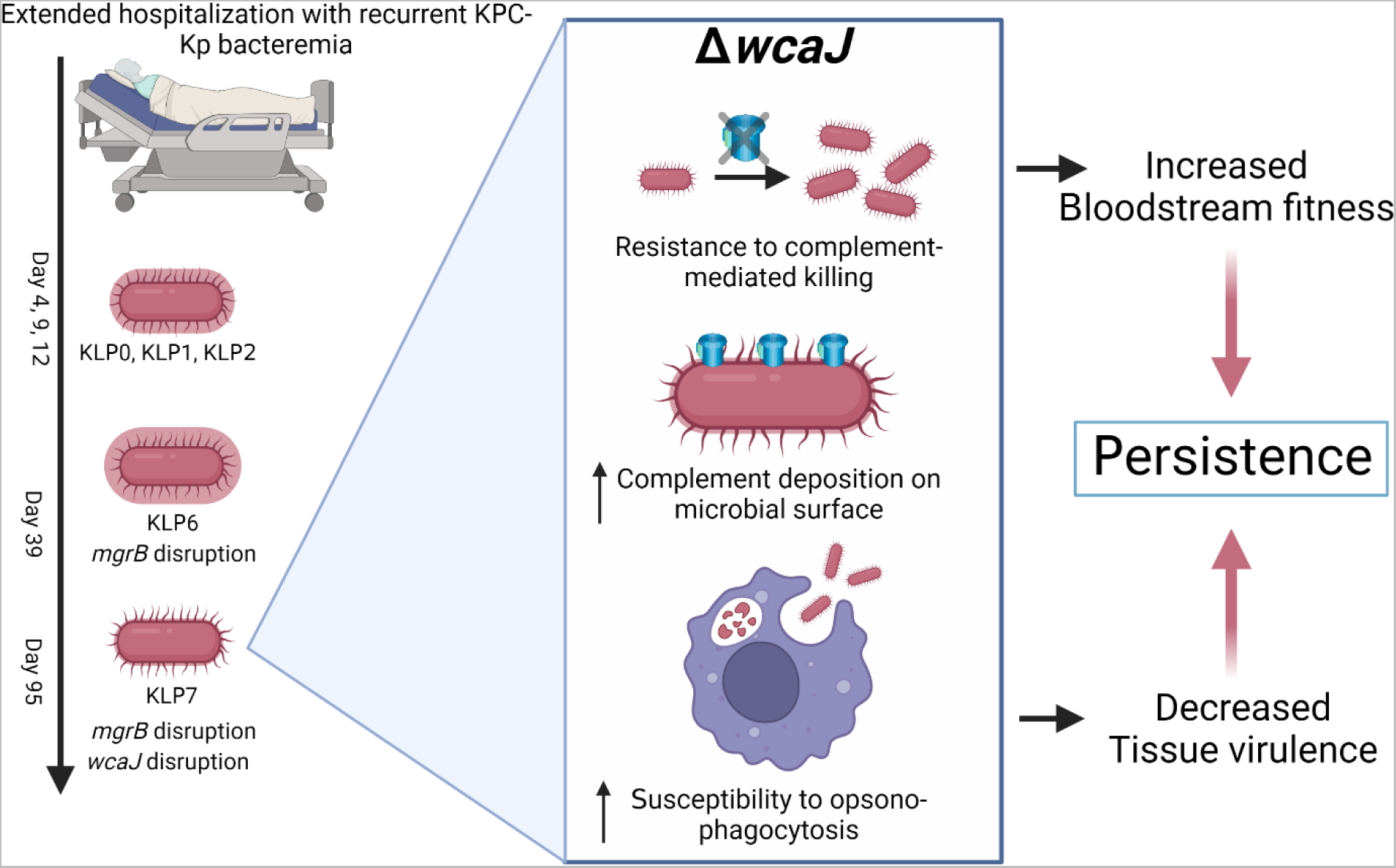
Graphical abstract. Created with BioRender.com

## Introduction

The expanding antibiotic resistance profile of *Klebsiella pneumoniae* is an increasing public health concern.(1–3) *Klebsiella pneumoniae* carbapenemase-producing *Klebsiella pneumoniae* (KPC-Kp) strains demonstrate extensive antimicrobial resistance and leave few therapeutic options during infection. This is of particular concern as KPC-Kp infections have been associated with high mortality rates in several published series.(4–6) However, KPC-Kp clinical strains, including the epidemic sequence type 258 (ST258), generally lack classical virulence determinants such as hypermucoviscous capsules belonging to the K1 and K2 serotypes. Furthermore, ST258 strains generally do not induce lethality in immunocompetent mice due to rapid clearance of the bacteria.(7–11) This paradox of KPC-Kp infection, where a relatively avirulent strain causes high morbidity and mortality in hospital settings, suggests deficiencies within the host defense program, which may provide fertile ground for evolving KPC pathogenicity. This is in contrast to hypervirulent strains, originally reported in East Asia in the 1980s, that cause invasive community-acquired infections associated with liver abscesses, necrotizing pneumonia, endophthalmitis, and meningitis and tend to be characterized by a hypermucoviscous phenotype.(1, 12–14) Therefore, there is likely an interplay of pathogen and host factors that influences the pathogenic potential of KPC-Kp strains during infection.(15)

A major mechanism of Kp pathogenic potential is the expression of a polysaccharide capsule that enables the bacteria to evade complement-mediated killing and phagocytosis(16, 17) and increases Kp pathogenicity in mice and humans.(16, 18–20) While the complement system can mediate direct serum killing of Gram-negative bacteria through membrane lysis as well as opsonization to enhance phagocytosis(1), there are widely varying reports of the susceptibility of KPC-Kp strains to serum-mediated killing.(8, 18, 19, 21) Here we sought to determine the mechanism of resistance of a series of clinical KPC-Kp isolates to serum-mediated killing to better understand the role of complement in the interaction between KPC-Kp and the host. We identified the *in vivo* development of complement resistance in serial KPC-Kp bloodstream isolates from a single patient with recurrent KPC-Kp bacteremia during an extended critical illness hospitalization. We identified a mobile genetic element insertion causing loss of function of the *wcaJ* gene that increased fitness in the bloodstream by providing resistance to complement-mediated serum killing. The same *wcaJ* mutation increased complement deposition on the microbial surface and enhanced opsono-phagocytosis, providing a mechanistic explanation for limited virulence and increased persistence in the host. These findings highlight a unique mechanism by which microbial evolution can occur within the host to simultaneously enhance bloodstream survival and limit virulence to promote longer term persistence.

## Results

### ST258 KPC-Kp clinical isolates demonstrate variable growth in pooled serum from healthy volunteers

Although ST258 KPC-Kp clinical isolates are generally thought to be sensitive to serum-mediated killing,(8, 19, 22) others have suggested that many KPC-Kp clinical isolates can persist in healthy human serum.(21) We conducted a screening assay for bacterial growth in serum using 59 ST258 clinical isolates from three separate academic medical centers (Fig 1A). We noted that 43 of the 59 isolates (73%) demonstrated little or no growth in serum, indicating serum sensitivity (Fig 1B). In contrast, 10/59 (17%) isolates demonstrated moderate growth, defined as between 2-fold to 5-fold increase in OD_600_ at 4 hours compared to baseline. An additional 6/59 (10%) isolates demonstrated high growth in serum, with a greater than 5-fold increase in OD_600_. Interestingly, we identified five serial KPC-Kp blood culture isolates with varying serum resistance profiles obtained from a single critically ill patient with prolonged hospitalization. We pursued further investigation of these isolates to better understand factors that contribute to serum resistance in KPC-Kp.

**Figure 1.**
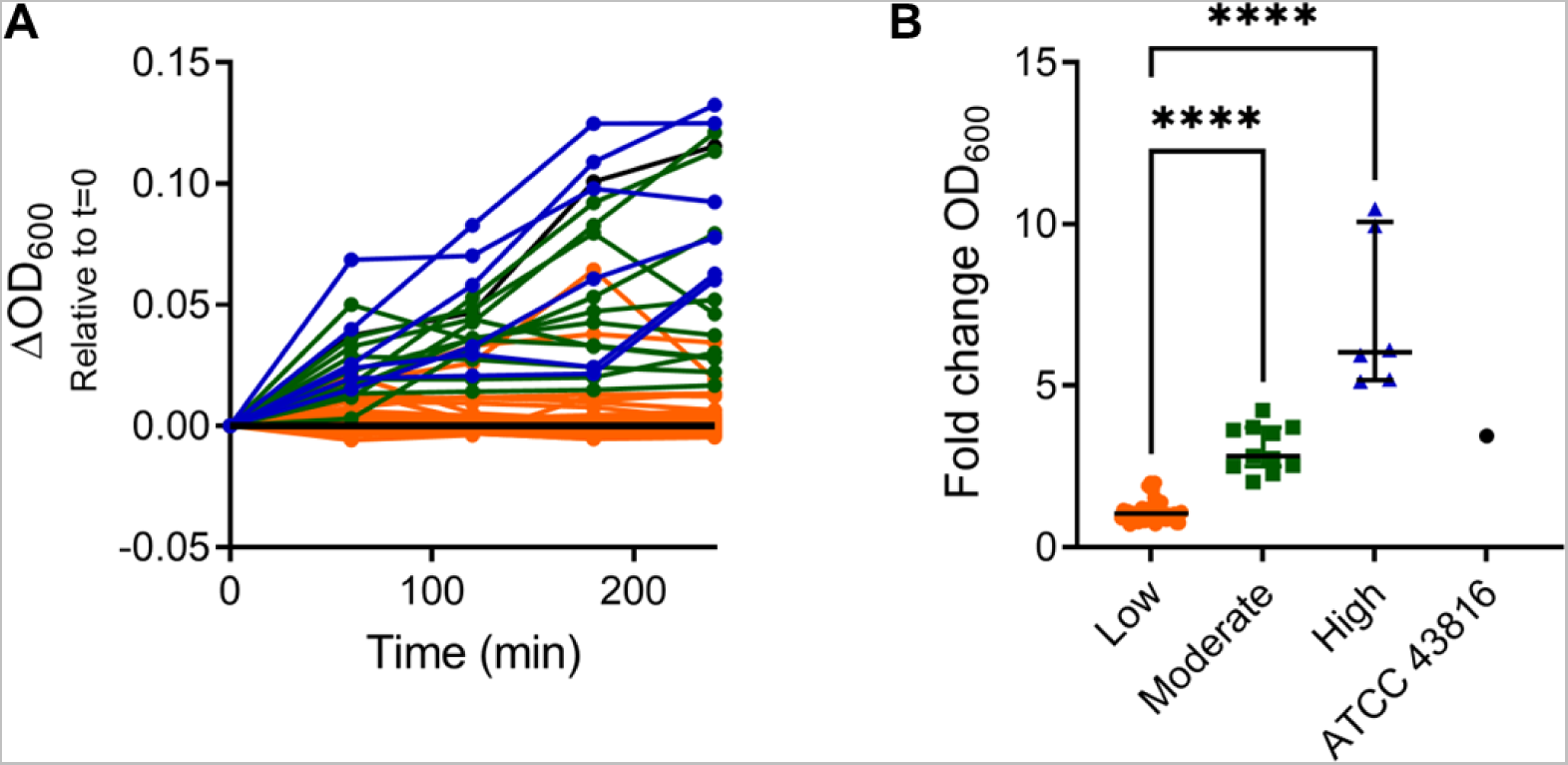
Sequence type 258 KPC-Kp clinical isolates demonstrate variable growth in pooled serum from healthy volunteers. (A) Optical density 600 nm of 59 clinical sequence type 258 (ST258) KPC-Kp isolates obtained from various loci of infection at three USA academic medical centers in the presence of pooled serum from healthy volunteers. Change in OD600 relative to time=0 during a single replicate screening trial is displayed. (B) Relative serum resistance of KPC-Kp strains were stratified into high (n=6, 10%), moderate (n=10, 17%), and low or no serum resistance (n=43, 73%) categories by fold change in OD600 at 4 hours relative to time 0. ATCC 43816 strain is displayed for reference. Each point represents a single clinical isolate, and the horizontal lines represent median and inter-quartile range. Kruskal-Wallis test with Dunn’s post hoc for comparison to “Low” group are displayed. ****p<0.0001 for post-hoc analysis.

### Patient Case History

The patient presented to the hospital for abdominal pain and jaundice and the medical history was notable for a liver transplant ten years prior that was managed with maintenance immune suppression. The patient had an extended acute care hospitalization (115 days) in both intensive care and long-term acute care units that was notable for persistent biloma and recurrent bloodstream infections with KPC-Kp. KPC-Kp was also isolated from hepatic fluid from the persistent biloma. The initial bloodstream KPC-Kp isolate was collected on hospital day 7, leading to the initiation of therapy with intravenous colistin, which was continued through discharge. The patient ceased to breathe 12 days after discharge, although the cause of death was not available to study investigators.

### In vivo development of serum and polymyxin resistance

We characterized five serial KPC-Kp blood culture isolates by whole-genome sequencing on the Illumina platform, which revealed that all five isolates – KLP0, KLP1, KLP2, KLP6, and KLP7 – were genetically related to one another (Fig 2A). KLP1 was also long-read sequenced on the PacBio platform and was thereafter used as the reference isolate. All five isolates were predicted to belong to ST258 clade I and encoded KL106/wzi29-type capsule loci. We identified three mutations with functional consequences in KLP2, which was isolated on hospital day 12, or five days after initiation of colistin treatment. These mutations were: 1) a T insertion into a polyT tract leading to a frame-shift mutation in the 3-oxosteroid 1-dehydrogenase gene *ksdD* (involved in lipid metabolism), 2) a 725-bp deletion (truncation) in the *cadC* transcriptional regulator gene of the *cadAB* operon,(23) and 3) a non-synonymous SNP, S132P, in the peptidyl-dipeptidase *dpc*.(24) On hospital day 39, KLP6 was isolated and was found to further contain an IS*5* family transposase insertion disrupting *mgrB*, which is the most common cause of colistin resistance in *K. pneumoniae* resulting in a 4-amino-4-l-arabinose modification of lipid A.(25–28) On hospital day 95, KLP7 was isolated from the patient’s bloodstream. This clinical isolate had a disruption in the capsule locus, with an IS*5* transposase insertion in the *wcaJ* gene.

**Figure 2.**
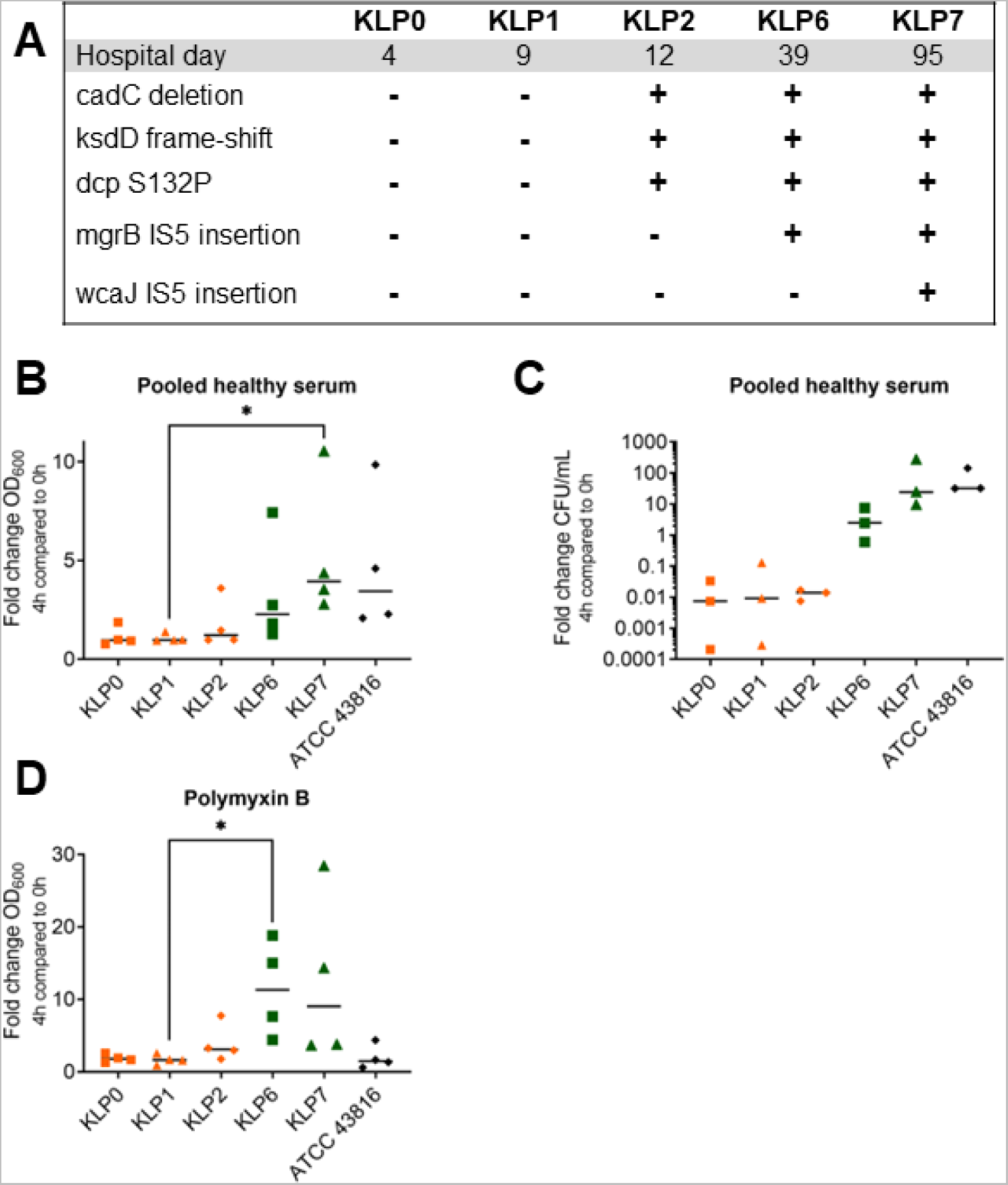
***In vivo* evolution of serum and polymyxin resistance in genetically-related serial ST258 KPC-Kp clinical isolates from the blood of a critically-ill patient treated with colistin.** Five genetically-related serial ST258 KPC-Kp isolates were obtained from a single patient with recurrent *K. pneumoniae* bacteremia who was treated with intravenous colistin during a 115-day hospitalization. (A) Temporal accumulation of mutations. Colistin treatment was initiated between collection of KLP0 and KLP1. Serial isolates were grown in serum and growth was quantified by (B) OD600 (N=4 trials) and (C) CFU (N=3 trials) after four hours incubation at 37°C. Results showing fold change at 4 hours compared to inoculum at 0 hours are displayed. (D) Serial isolates were also grown in polymyxin B (85% vol/vol) and growth was quantified by OD600 at 4 hours and compared to 0 hour baseline. ATCC 43816, a K2 serotype *K. pneumoniae* research strain, is displayed for reference. Statistics by Friedman test with Dunn’s post-hoc correction for multiple comparisons. *Post-hoc p<0.05 for KLP7 compared to KLP1 in Panel B and KLP6 compared to KLP1 in Panel D, no other comparisons were statistically significant.

To assess the functional consequences of the genetic mutations that accumulated during extended hospitalization and recurrent bloodstream infections in this patient, we first repeated testing of the serum resistance profiles of the five serial isolates (Fig 2B). As anticipated, we found increased serum resistance in KLP6 and KLP7, which was validated by CFU growth in serum at 4 hours compared to control (Fig 2C) . Notably, KLP7 reached statistically significant increased growth as measured by fold change at 4 hours from baseline in pooled healthy serum compared to KLP1 and was comparable to the well documented serum-resistant ATCC 43816 reference strain.(19, 29) We further documented substantial growth of KLP6 (p<0.05 compared to KLP1) and KLP7 in 85% (v/v) polymyxin B (Fig 2D), which is consistent with disruption of *mgrB*.(25) These findings suggested that extended colistin treatment and recurrent bloodstream infections led to selection of *mgrB* and *wcaJ* variants. Therefore, we sought to further phenotype these serial bloodstream isolates to determine how they were able to escape both colistin and serum-mediated killing.

### wcaJ mutation is associated with moderate biofilm production, impaired osmotic tolerance, and capsule alteration

As the patient was managed with several indwelling venous, arterial, and percutaneous catheters during extended critical illness, we examined the biofilm production potential of the five serial isolates using a crystal violet assay. We found significantly increased biofilm production in the KLP2 isolate (Fig 3A), which was isolated from the blood approximately 8 days after placement of a central venous catheter. Notably, all venous and arterial catheters were removed 6 days later on hospital day 18 due to persistent Kp bacteremia on hospital days 15-17. No central venous catheter was placed until a temporary hemodialysis catheter was placed on hospital day 23. We were curious if the accumulation of genetic mutations in these isolates altered their susceptibility to abnormal growth conditions. We found significantly decreased growth in KLP7 compared to KLP1 when grown in tryptic soy broth with 50% vol/vol sterile de-ionized water (Fig 3B), suggesting decreased osmotic tolerance in the isolate with a *wcaJ* mutation. In contrast, we found no differences in growth of serial isolates in tryptic soy broth with and without sodium chloride supplementation (2.5% and 5% weight/volume, data not shown). We then assessed the role of accumulated genetic mutations in capsule formation, and found decreased uronic acid production(7) in KLP7 isolate (Fig 3C), which is consistent with the predicted role of the *wcaJ* glycosyltransferase gene in capsule formation. When streaked on BHI agar supplemented with sucrose and Congo red,(30) we noted gross differences in the appearance and size of colonies when comparing KLP1 with the KLP6 and KLP7 isolates (Fig 3D). These gross changes were confirmed by transmission electron microscopy with ruthenium red staining (Fig 3E) as the KLP7 isolate demonstrated marked reduction in polysaccharide capsule on the bacterial surface. Together, these data suggest that progressive genetic modifications identified in serial KPC-Kp bloodstream isolates conferred serum resistance through alterations in the capsular polysaccharide.

**Figure 3.**
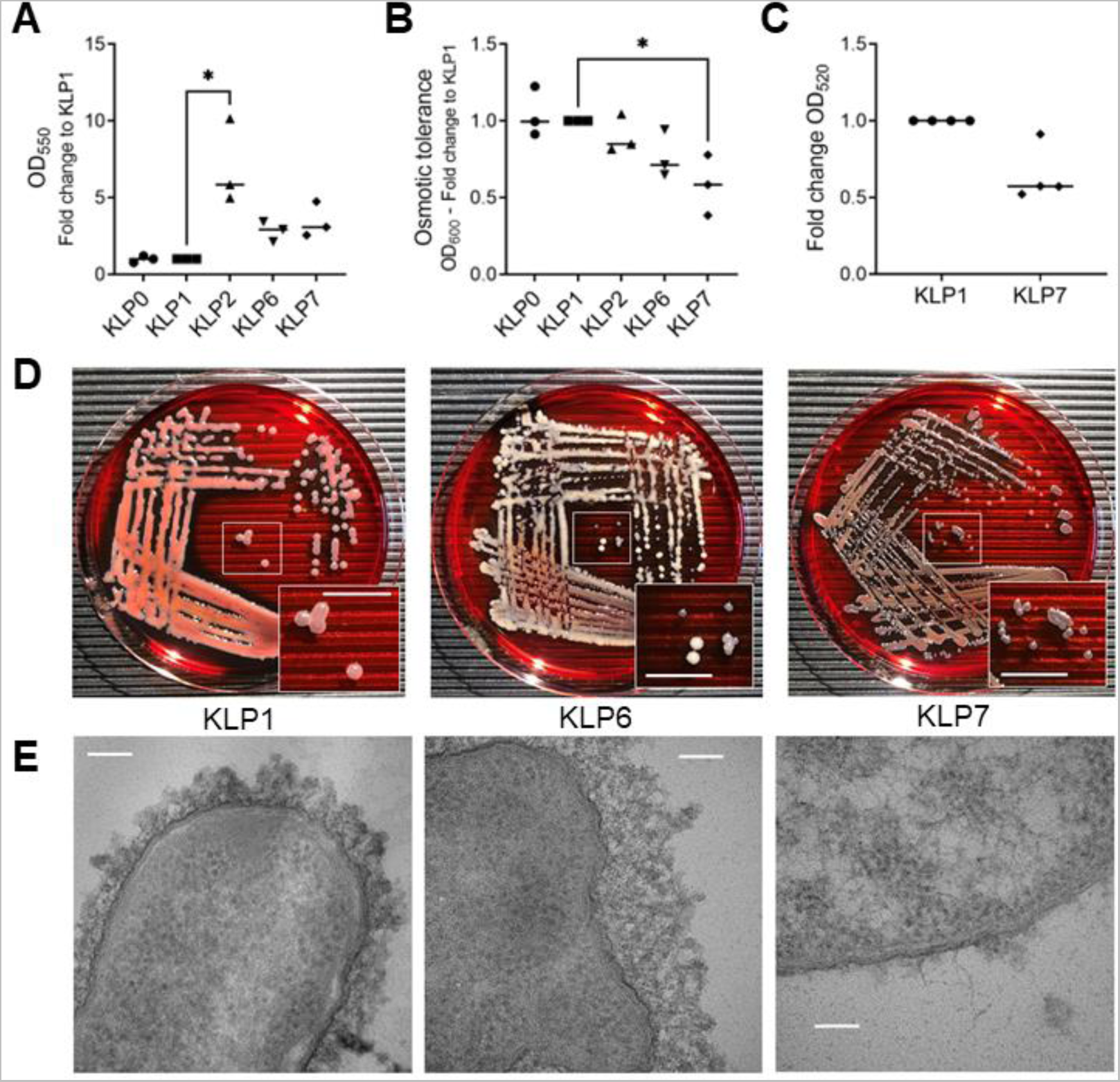
A clinical KPC-Kp isolate with *wcaJ* disruption exhibits moderate biofilm production, impaired osmotic tolerance, and capsule alteration. Five serial ST258 KPC-Kp blood isolates obtained from a single patient with persistent *K. pneumoniae* bacteremia demonstrate (A) differences in biofilm formation as quantified by crystal violet assay. The median result of three separate trials by fold change in optical density 550 nm compared to the genetic reference strain KLP1 are displayed. (B) Changes in osmotic tolerance as measured by fold change in optical density at 600 nm during incubation in tryptic soy broth with 50% sterile de-ionized water at 4 hours. The median result of three separate trials quantifying fold change to the genetic reference strain (KLP1) are displayed. (C) Uronic acid content of capsular polysaccharide extract as measured by fold change in optical density 520 nm. The median result of four separate trials quantifying fold change to the genetic reference strain (KLP1) are displayed. (D) Gross appearance of KLP1, KLP6, and KLP7 isolates grown on brain heart infusion agar supplemented with 5% (weight/volume) sucrose and Congo red. White boxes are displayed in insets and white scale bar indicates approximately 10 mm. (E) Transmission electron microscopy of isolates KLP1, KLP6, and KLP7 after ruthenium red staining. Representative images are displayed at 200,000X magnification. White scale bar indicates 100 nm. *p<0.05 for Panel A (KLP2 compared to KLP1) and Panel B (KLP7 compared to KLP1) by Kruskal-Wallis Test with Dunn’s post-hoc for multiple comparisons.

### wcaJ mutant clinical KPC-Kp isolate exhibits resistance to complement-mediated serum killing and increased C3 and membrane attack complex binding

Because the complement system is a crucial component of host defense against Kp in the blood, we examined whether the *wcaJ* mutation in KLP7 conferred serum resistance through evasion of complement-mediated killing. Normal healthy serum robustly killed the KLP1 isolate (Fig 4A), although killing was significantly impaired in the presence of serum depleted of complement component C3. Notably, mixing serum depleted of C3 in 1:1 fashion with healthy serum restored effective killing, confirming that complement is the primary component of serum-mediated killing of KLP1. In contrast, the KLP7 isolate was resistant to serum killing in normal healthy serum, C3-depleted serum, and mixed serum. We hypothesized that KLP7 resists serum killing by limiting complement binding.(31) Surprisingly, we found significantly increased binding of complement C3 (Fig 4B) and membrane attack complex (Fig 4C) to the KLP7 isolate compared to KLP1. We confirmed this unexpected finding by performing flow cytometry for C3 and membrane attack complex (MAC) binding to the KLP1, KLP6, and KLP7 isolates. We detected increased binding of C3 (Fig 4D) and MAC (Fig 4E) to the KLP7 isolate, which was absent after incubation with bacteria alone or with C3-depleted and C6-depleted serum, respectively. We next performed immuno-electron microscopy to confirm increased complement binding to the KLP7 isolate and noted extensive binding of C3 to the KLP7 surface that was largely absent with KLP1 (Fig 4F). Taken together, these orthogonal approaches confirmed increased complement binding to the serum-resistant KLP7 strain. We therefore sought to investigate whether the *wcaJ* gene mutation conferred this phenotype.

**Figure 4.**
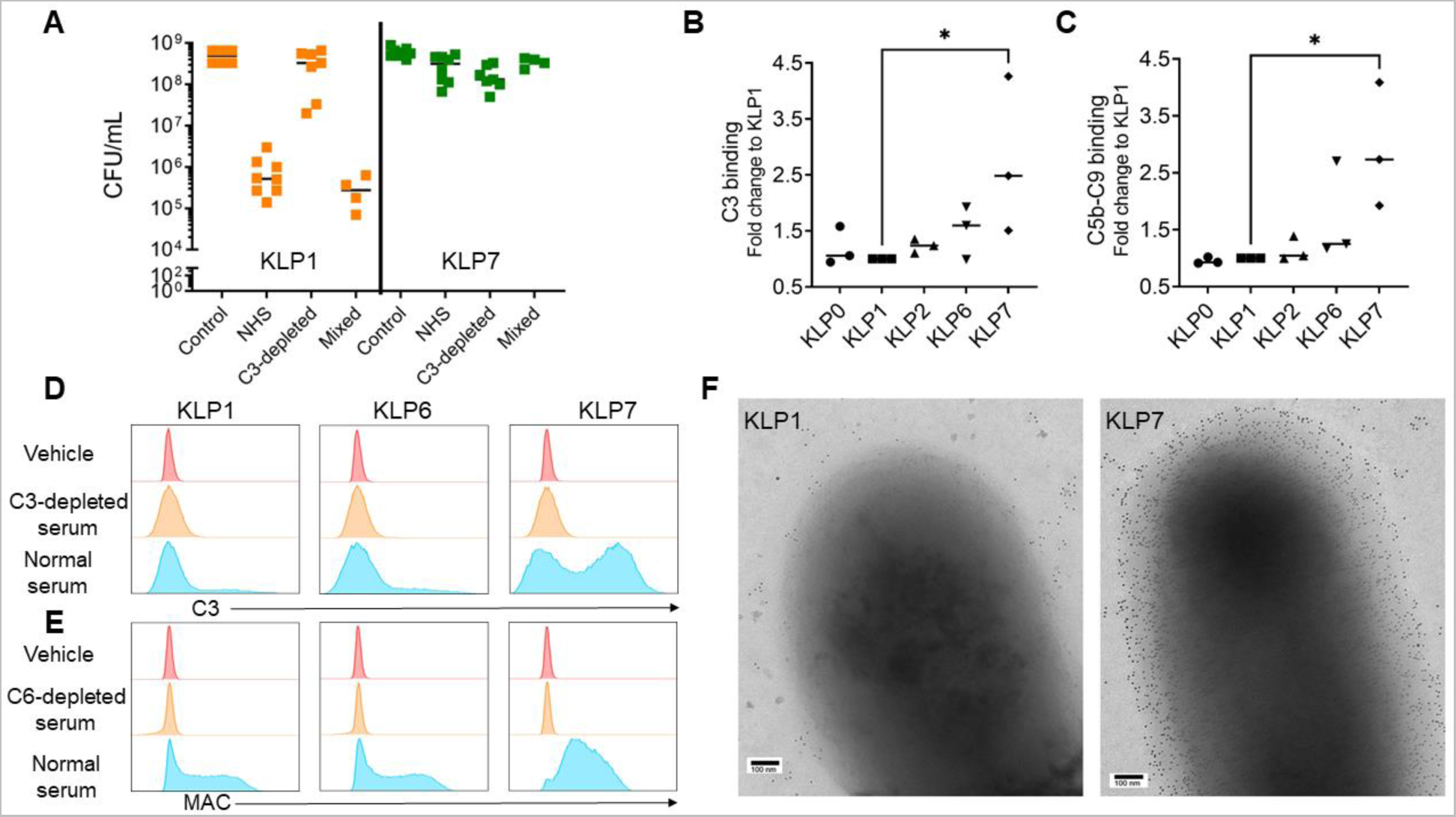
The *wcaJ* mutant KPC-Kp isolate exhibits resistance to complement-mediated serum killing despite increased binding by complement component C3 and the membrane attack complex. (A) Mixing study of KPC-Kp clinical isolates KLP1 (genetic reference) and KLP7 (IS*5* insertions in *mgrB* and *wcaJ*) after incubation for 4 hours in pooled serum from healthy volunteers and serum depleted of complement component C3. Groups include isolates grown in 85% vol/vol solution of sterile phosphate buffered saline (control; n=8 trials), healthy serum (NHS; n=8 trials), or sera depleted of complement component C3 (C3 depleted; n=7 trials). In select experiments, C3-depleted serum was also mixed 1:1 with healthy serum (n = 4 trials). Each point represents the median colony-forming units (CFU) per mL result from a single trial. (B-C) Direct ELISA of five serial ST258 KPC-Kp isolates to quantify binding of complement components C3 and C5b-C9 (membrane attack complex) after incubation with diluted pooled healthy serum for 30 minutes. The result of three separate trials quantifying fold change to the KLP1 reference isolate with median result are displayed. *p<0.05 by Kruskal-Wallis Test with Dunn’s post-hoc. (D-E) Flow cytometry histograms of D) C3b and E) MAC binding for KLP1, KLP6 (*mgrB* insertional inactivation), and KLP7 (*mgrB* and *wcaJ* insertional inactivation) after incubation for 30 minutes with diluted pooled healthy serum. (F) Immunogold transmission electron microscopy of KLP1 and KLP7 incubated for 30 minutes with diluted pooled healthy serum and primary antibody for C3 followed by colloidal gold secondary antibody. Representative images at 100,000X magnification are displayed.

### Deletion of wcaJ increases complement binding and opsono-phagocytosis

To confirm the role of *wcaJ* in altered complement binding, we deleted *wcaJ* in the KLP1 isolate background to generate an isogenic pair of wild-type and Δ*wcaJ* strains (Supplemental Figure 1). We noted similarity in colony morphology between KLP1Δ*wcaJ* and KLP7 when streaked onto BHI agar supplemented with sucrose and Congo red (Fig 5A). Scanning electron microscopy also confirmed marked changes in capsule abundance and architecture in KLP1Δ*wcaJ* that resembled KLP7 (Fig 5B). Similar to KLP7, KLP1Δ*wcaJ* was resistant to serum killing in normal healthy serum, C3-depleted serum, and mixed serum (Fig 5C). To confirm that deletion of *wcaJ* mediates resistance to complement-mediated serum killing, we performed complementation of *wcaJ* into the KLP1 Δ*wcaJ* strain, which partially restored serum sensitivity (Fig 5D) thereby confirming that deletion of *wcaJ* mediates serum resistance. Furthermore, flow cytometry confirmed increased binding of complement C3 to KLP1Δ*wcaJ* when compared to KLP1 (Fig. 5E).

**Figure 5.**
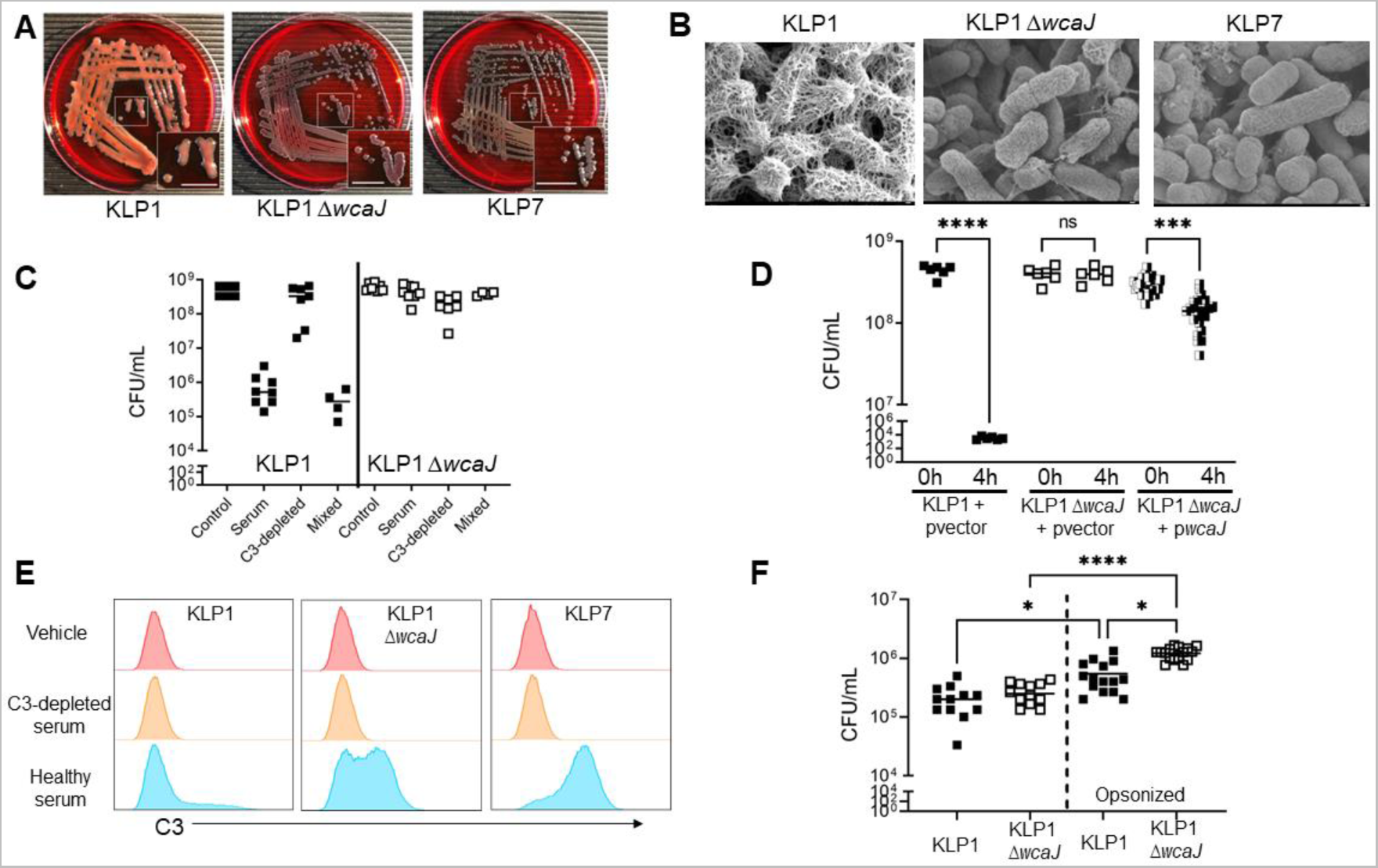
Deletion of *wcaJ* decreases complement killing but increases complement binding and opsono-phagocytosis. The *wcaJ* gene was deleted from the reference clinical isolate KLP1 by homologous recombination (KLP1*ΔwcaJ*). (A) Gross appearance of KLP1, KLP1Δ*wcaJ*, and KLP7 grown on brain heart infusion agar supplemented with sucrose and Congo red. White boxes are displayed in insets and white scale bar indicates approximately 10 mm in inset. (B) Scanning electron microscopy of isolates KLP1, KLP1Δ*wcaJ*, and KLP7. Representative images are displayed at 30,000X magnification. White scale bar in lower right corner indicates 100 nm. (C) Mixing study of KLP1 and KLP1Δ*wcaJ* in pooled serum from healthy volunteers and serum depleted of complement component C3 after incubation for 4 hours. Groups include isolates grown in 85% vol/vol solution of sterile phosphate buffered saline (control; n=8 trials), healthy serum (NHS; n=8 trials), or sera depleted of complement component C3 (C3 depleted; n=7 trials). In select experiments, C3 depleted serum was also mixed 1:1 with healthy serum (n=4 trials). Each point represents colony-forming units (CFU) per mL from a single trial. (D) Comparison of CFU/mL at 0 hours and after 4 hours in pooled healthy serum in KLP1 and KLP1Δ*wcaJ* with empty plasmid vector (pvector) or KLP1Δ*wcaJ* with plasmid containing *wcaJ* (p*wcaJ*) that partially restores serum sensitivity. Each point represents a technical replicate from a single representative trial (N=2 biological replicates for KLP1 and KLP1Δ*wcaJ* with empty plasmid vector; N=8 biological replicates for KLP1Δ*wcaJ* with plasmid complementation of *wcaJ*). (E) Flow cytometry histograms of MAC binding for KLP1, KLP1Δ*wcaJ*, and KLP7 after incubation for 30 minutes with diluted pooled healthy serum. (F) KLP1 and KLP1Δ*wcaJ* were synchronously incubated with RAW 264.7 macrophage cells in culture. In select experiments, bacteria were opsonized in 20% vol/vol human serum on ice. After 60 minutes, cells were incubated with polymyxin B for 30 minutes to kill extracellular bacteria then cells were lysed and intracellular bacteria were counted after serial dilution and overnight incubation on tryptic soy agar. Each point represents a single replicate from multiple trials (n=3 trials without opsonization, n=4 trial with opsonization). Statistics by Kruskall-Wallis with Dunn’s post-hoc in Panels D and F – post-hoc significance is displayed. *p<0.05, ***p<0.001, ****p<0.0001.

If loss of *wcaJ* function mediates resistance to complement-mediated serum killing, we wondered why this increased virulence did not lead to overwhelming infection and tissue destruction in this patient. Given the increased complement deposition observed on the microbial surface of KLP1Δ*wcaJ*, we hypothesized that the KLP1Δ*wcaJ* mutant might have been rendered more susceptible to opsono-phagocytosis, thereby limiting its pathogenicity. In the absence of pre-incubation with serum, there was no significant difference in phagocytosis of KLP1Δ*wcaJ* by RAW264.7 macrophages compared to the KLP1 parent strain (Fig 5E). In contrast, opsonization with 20% serum increased phagocytic uptake of KLP1Δ*wcaJ* when compared to opsonized KLP1 and non-opsonized KLP1Δ*wcaJ* (Fig 5E). These results indicate that *wcaJ* loss-of-function mutation increases resistance to complement-mediated killing, but also increases complement binding and susceptibility to opsono-phagocytosis. We therefore sought to test the *in vivo* role of opsono-phagocytosis during KLP1Δ*wcaJ* infection.

### Disabling opsono-phagocytosis through depletion of alveolar macrophages impairs in vivo control of KPC-Kp ΔwcaJ in an acute lung infection model

As *wcaJ* loss-of-function rendered KLP1 more susceptible to opsono-phagocytosis, we tested the *in vivo* impact of disabling opsono-phagocytosis on the pathogenicity of the KLP1Δ*wcaJ* mutant. We depleted alveolar macrophages in wildtype C57Bl/6J mice via intra-tracheal administration of clodronate liposomes 24 hours prior to intra-tracheal infection with KLP1Δ*wcaJ* and performed necropsy 24 hours post-infection (Fig. 6A). Flow cytometry of BALF (Supplemental Figure 2) collected at necropsy confirmed a nearly 50-fold decrease in alveolar macrophages with clodronate treatment (Fig 6B) but no significant difference in neutrophil influx into the airspaces compared to liposome vehicle (Fig 6C). To determine whether fluid in the alveolar space was sufficient to opsonize KLP1Δ*wcaJ*, we performed an *ex vivo* opsonization assay. We found that BALF from mice inoculated with KLP1Δ*wcaJ* was sufficient to enable C3 binding, as determined by direct ELISA (Fig 6D). This finding supports the concept that pathogens can be opsonized in the alveolar space during infection, which is consistent with prior reports(32). We observed increased BALF protein content in mice treated with clodronate (Fig 6E), which is a non-specific marker of increased lung microvascular permeability. Most importantly, we found that mice depleted of alveolar macrophages by clodronate treatment exhibited impaired control of KLP1Δ*wcaJ* as determined by significantly increased CFU counts in lung homogenate (Fig 6F). These data suggest that the KLP7 isolate bearing a *wcaJ* mutation is more readily phagocytosed by tissue-resident immune cells, thereby limiting its virulence.

**Figure 6.**
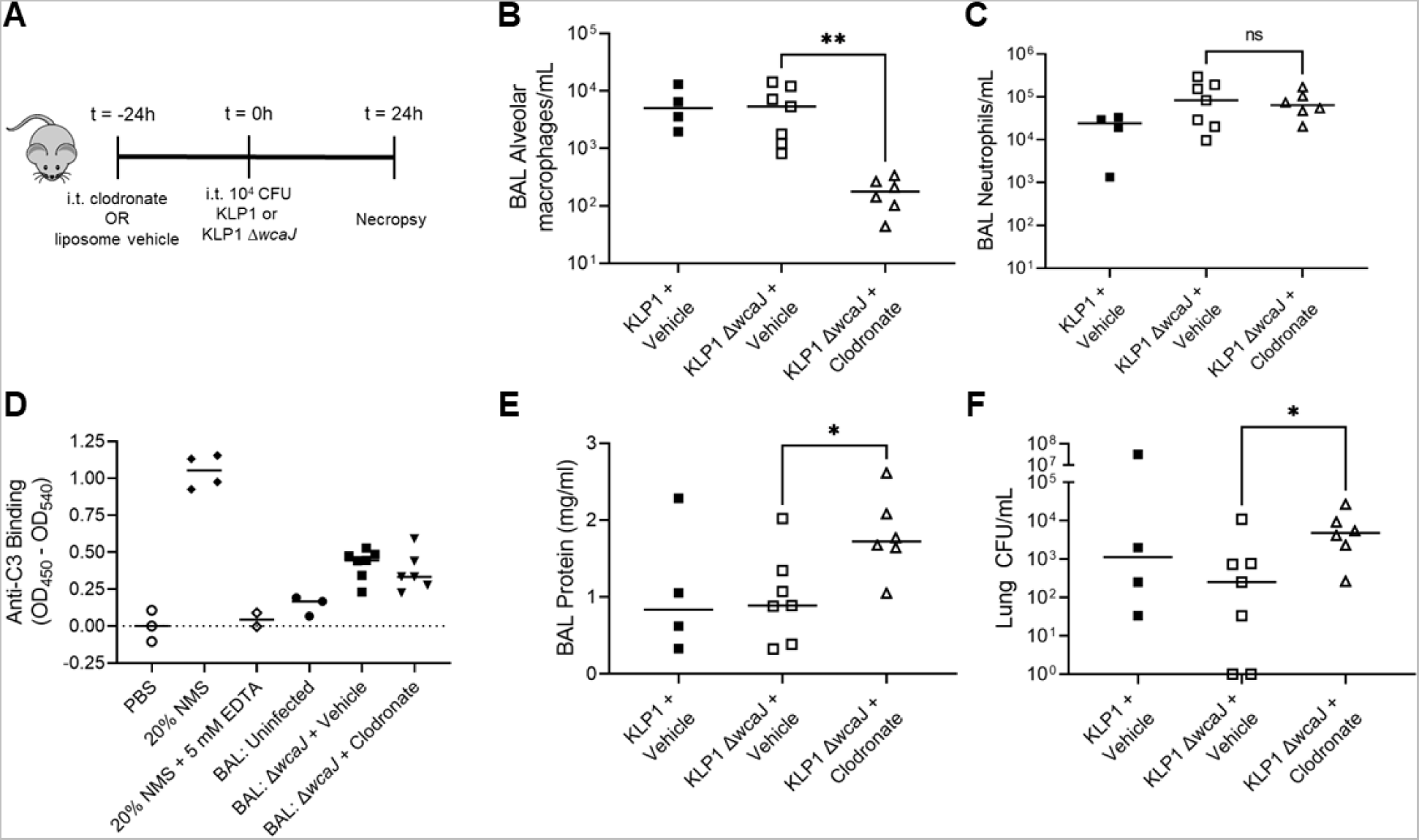
Disabling opsono-phagocytosis through depletion of alveolar macrophages impairs *in vivo* control of KPC-Kp with *wcaJ* deletion in an acute lung infection model. (A) Experimental schematic demonstrating that wildtype C57Bl/6J mice were intra-tracheally treated with liposome control (n=11) or clodronate (n=6) to deplete alveolar macrophages followed 24 hours later by intra-tracheal inoculation with 10^4^ colony forming units (CFU) of KLP1 (n=4 after vehicle) or KLP1 Δ*wcaJ* (n=7 after vehicle; n=6 after clodronate) with subsequent necropsy at 24 hours post-infection. Flow cytometry was performed on broncho-alveolar lavage (BAL) cells obtained at necropsy with (B) alveolar macrophage and (C) neutrophil counts per milliliter quantified by counting beads confirming macrophage depletion by clodronate treatment. (D) *Ex vivo* BAL opsonization assay using direct ELISA (optical density 450 nm with subtraction of background optical density 540 nm) of anti-mouse-C3 binding to KLP1 Δ*wcaJ* after 1 hour incubation with phosphate buffered saline (PBS, median value was utilized to determine background binding, n=3), 20% v/v normal mouse serum (NMS, n=4) with or without EDTA (n=2), and BAL from uninfected mice (n=3) or BAL obtained at necropsy from experimental animals. (E) BAL protein was quantified by BCA assay. (F) Lung CFU per mL. Each point represents a single mouse. **p<0.01 and *p<0.05 by Mann Whitney test. ”ns” = non-significant. The results of KLP1 after vehicle treatment group are displayed for reference.

### Compstatin-mediated inhibition of complement C3 impairs opsono-phagocytosis of KPC-Kp with wcaJ deletion in human whole blood

Complement component C3 is proteolytically cleaved into C3a and C3b, the latter of which is a key opsonin used by several phagocytic receptors. Therefore, we tested whether compstatin, a specific inhibitor of human C3 cleavage and activation, would reduce opsono-phagocytosis of KPC-Kp strains. After pHrodo labeling of heat-killed KLP1, KLP1Δ*wcaJ*, KLP6, and KLP7, we performed a flow cytometry-based assay of opsono-phagocytosis 30 minutes after inoculation into lepirudin-treated human whole blood with or without compstatin.(33, 34) As described by others, we utilized forward and side scatter gating to identify granulocytes and monocytes in whole blood (Figure 7A).(33) Compstatin treatment impaired opsono-phagocytosis of KLP1Δ*wcaJ* and KLP7 (Figure 7B, representative trial with granulocytes) as demonstrated by significantly decreased percentages of granulocytes and monocytes positive for pHrodo staining in compstatin-treated whole blood compared to vehicle (PBS) treatment (Figure 7C, overall p<0.01 with post-hoc significance p<0.05 for KLP1Δ*wcaJ* compared to KLP1 and for KLP7 compared to KLP1 and KLP6). Therefore, complement-mediated opsonization of KPC-Kp with *wcaJ* deletion promotes susceptibility to opsono-phagocytosis, despite resistance to complement-mediated killing. Taken together, these data suggest that opsono-phagocytosis is a crucial component of host defense against KPC-Kp, and that *wcaJ* deletion limits pathogenicity and likely contributes to persistent infection.

**Figure 7.**
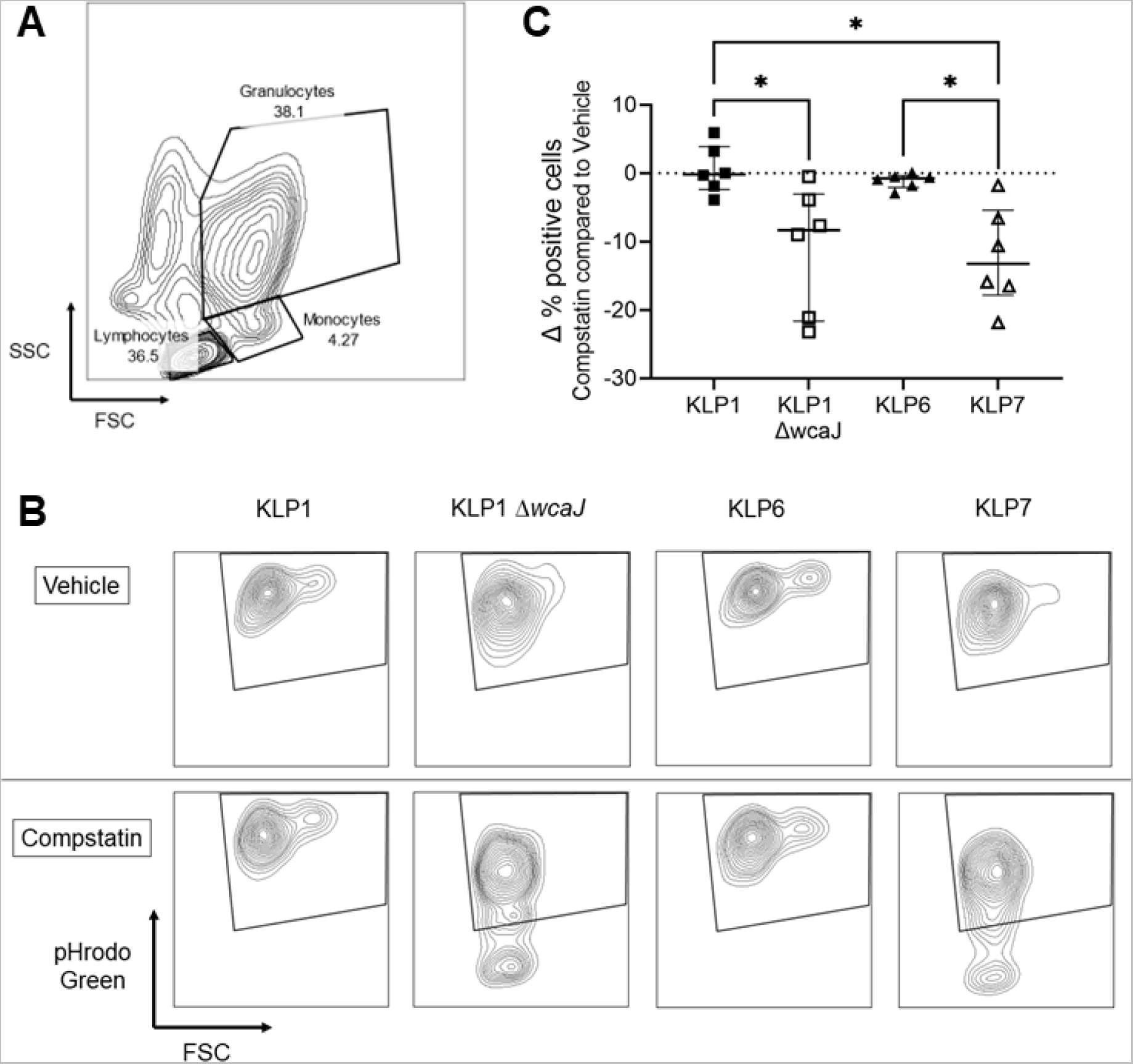
Compstatin-mediated inhibition of complement C3 impairs opsono-phagocytosis of KPC-Kp with *wcaJ* deletion in human whole blood. Flow cytometry was performed on fresh human whole blood treated with 50 µg/mL lepiriduin (N=3 separate trials with blood from a single human donor). (A) Representative gating with forward scatter (FSC, X-axis) and side scatter (SSC, Y-axis) identification of granulocytes, monocytes, and lymphocytes. (B) Representative result of pHrodo green (Y-axis) measurement of opsono-phagocytosis at 30 minutes by granulocytes for each bacterial isolate from a single trial. The trapezoidal box indicates pHrodo green positive cells. The top row received vehicle (PBS) treatment and the bottom row received 167 µM compstatin treatment demonstrating decreased pHrodo green positive granulocytes with compstatin treatment with KLP1 Δ*wcaJ* and KLP7 isolates. (C) Summary results of change in % positive cells with compstatin treatment in granulocyte and monocyte gates from all trials. Each point represents a single gate. Statistics by Kruskal-Wallis Test with Dunn’s post-hoc for multiple comparisons. Significant post-hoc p-values are displayed.

## Discussion

Our findings provide a unique assessment of host-pathogen interaction and bacterial evolution within a single critically ill patient marked by recurrent KPC-Kp bloodstream infection. We note that the patient survived numerous episodes of KPC-Kp bloodstream infection, with the infecting strain of KPC-Kp evolving several mutations over time. The final bloodstream isolate exhibited a loss-of-function mutation in the *wcaJ* gene that severely limited polysaccharide capsule biosynthesis and conferred resistance to complement-mediated serum killing. While this loss-of-function mutation in the *wcaJ* gene increased the fitness of the bacterium in the bloodstream, the same mutation also increased complement deposition on the microbial surface, which increased susceptibility to opsono-phagocytosis by the host. These findings suggest that the functional consequences of *wcaJ* mutation were both enhanced bloodstream fitness and limited tissue pathogenicity, which ultimately promoted bacterial persistence within the host blood.

*K. pneumoniae* extracellular capsule is a key virulence factor and modulation of genes critical to the polysaccharide capsule appears to be a major component of KPC-Kp evolution and fitness in the human host.(7, 35–37) The *wcaJ* gene resides within the *cps* gene cluster and encodes an enzyme that attaches the first glucose-1-phosphate sugar of the growing exopolysaccharide capsule onto undecaprenyl phosphate in the cytoplasm near the inner membrane.(38) Others have demonstrated that deletion of *wcaJ* in the K1 serotype SGH10 strain can promote gut persistence.(36, 37) Furthermore, it has recently been demonstrated that capsule deficient KPC-Kp isolates exhibit increased persistence in the host with the urinary tract as a common reservoir.(7) Notably, insertion sequences conferring loss-of-function in *wbaP*, a homolog of *wcaJ,* were the most commonly identified genetic modifications.(7) Still others have suggested that genetic deletion of *wcaJ* from the K3 serotype reference research strain ATCC 13883 contributes to increased biofilm formation and modulation of phagocytosis that may support persistence compared to the parent strain.(39) Therefore, capsule deficiency mediated by *wcaJ* loss-of-function appears to contribute to persistence of *K. pneumoniae* strains.

Here, we utilize a unique clinical circumstance of recurrent KPC-Kp bacteremia to understand why capsule biosynthesis loss-of-function may support persistence. We found that loss-of-function in the *wcaJ* gene establishes a bloodstream niche for KPC-Kp through evasion of complement-mediated killing. That *wcaJ* loss-of-function promotes a bloodstream niche for KPC-Kp is supported by a prior report of serial mutations that developed during the course of 4.5 years of abdominal carriage of KPC-Kp in a patient.(40) Notably, *in vivo* development of *wcaJ* mutation was documented in a bloodstream clinical isolate obtained during periods of cholangitis.(40) In fact, the *wcaJ* mutant was the only KPC-Kp isolate of 18 clinical isolates to be collected from the bloodstream.(40) We can only speculate as to the site of origin that led to KPC-Kp persistence in the patient described herein, but we note a clinical report of biloma at presentation that remained during the entire hospitalization and required a percutaneous trans-hepatic biliary drain.

We note the novel finding that a *wcaJ* loss-of-function mutation was sufficient to induce complement resistance in a KPC-Kp isolate. Capsular polysaccharide is a key pathway by which Kp strains resist complement-mediated killing(41, 42). Therefore, the complement resistance provided by *wcaJ* loss-of-function mutation was somewhat unexpected because the mutation demonstrated reduced capsule content, which is generally thought to increase susceptibility to complement mediated killing in *K. pneumoniae* strains(1, 29, 42). However, genetic diversity of the CPS gene island is a prominent characteristic of the ST258 lineage (35). and it is important to note that the biophysical properties and functional consequences of capsule modulation may be different between hypermucoviscous Kp strains and KPC-Kp isolates. For example, induction of colistin resistance can alter the biophysical characteristics of the Kp capsule, potentially hardening the capsule structure(43). Furthermore, others have demonstrated complex interactions between complement factors and Kp capsule properties in a series of serum-resistant, extended-spectrum β-lactamase-producing Kp bloodstream isolates(44). Therefore, although *wcaJ* mutation limits extra-capsular polysaccharide production, the functional consequences of the capsule alteration are increased resistance to complement-mediated killing through a yet to be determined mechanism, which may be of importance to other ST258 isolates.

Despite the complement resistance conferred by *wcaJ* loss-of-function, we were also surprised to find increased complement binding. Importantly, we note increased binding of the MAC, which is responsible for killing Gram-negative bacteria by forming pores in the bacterial membrane. Others have suggested that capsule-deficient non-ST258 Kp strains may resist killing by allowing complement deposition and activation, but sequestering this complement activity at a distance from the bacterial membrane to limit membrane damage(41). MAC binding in the absence of bacterial lysis has also been demonstrated in ESBL-producing Kp clinical blood stream isolates, although the authors did not describe remote binding of MAC, suggesting other potential mechanisms of resistance to MAC lysis(44). Notably, local action of complement C5 convertases at the bacterial membrane as well as direct anchoring of MAC precursors such as C5b-C7 may be crucial for execution of MAC pore formation and Gram-negative bacterial lysis(45, 46) although this has not been investigated in the ST258 lineage. Finally, it is possible that increased complement binding may lead to saturation of early MAC proteins leading to steric hindrance such that final assembly of the large multi-protein C9 complex that forms a β-barrel structured pore may be impaired(47, 48).

Although resistance to complement mediated killing is generally thought to confer virulence in *K. pneumoniae*, we noted limited tissue virulence with *wcaJ* loss-of-function. We propose that although the *wcaJ* mutation increased fitness in the bloodstream, the associated accentuation of complement binding also increased susceptibility to opsono-phagocytosis and limited tissue virulence. This finding may provide another perspective to understand how interaction between the host and KPC-Kp strains determines their pathogenic potential. We have previously shown that functional deficiency of the alternative complement pathway can increase host susceptibility to a KPC-Kp clinical isolate in serum-killing assays and mouse models of disseminated pneumonia(18). Yet, KPC-Kp isolates have a wide range of serum susceptibility and KPC-Kp infections are not uniformly fatal. In fact, the patient in our case was discharged to home 12 days after recovery of the complement-resistant *wcaJ* mutant isolate. Phagocytosis may serve as an additional layer of host defense to clear KPC-Kp bloodstream isolates that are complement-resistant. We further demonstrate that alveolar fluid alone was sufficient to opsonize the KLP1Δ*wcaJ* strain, that depletion of alveolar macrophages increased tissue pathogenicity of the *wcaJ* mutant, and that complement-mediated processes promote opsono-phagocytosis of *wcaJ* mutants in human whole blood. Therefore, opsono-phagocytosis is a host defense mechanism that counters the complement-resistance imparted by the *wcaJ* mutation. The implications of these findings for other KPC-Kp isolates are unclear, but the importance of complement and phagocytosis in defending the lung against Gram-negative pathogens has been shown repeatedly (18, 20, 49, 50).

In summary, we describe the *in vivo* development of serum resistance through the accumulation of genetic mutations in a patient with extended hospitalization and recurrent KPC-Kp bloodstream infection culminating in an IS*5* transposon insertion conferring loss of function mutation in the *wcaJ* gene. Despite conferring resistance to complement-mediated serum killing, the *wcaJ* mutation also led to increased complement binding and increased susceptibility to opsono-phagocytosis. Acquisition of these two traits led to persistence within the host by increasing bloodstream fitness and limiting tissue pathogenicity. Further work is needed to understand the wider clinical consequences of *wcaJ* loss-of-function in KPC-Kp clinical isolates, and to understand the molecular basis of persistence mutations that evolve within the host.

## Methods

### Bacteria

Clinical KPC-Kp isolates were obtained from three academic medical centers: University of Pittsburgh, University of Wisconsin, and University of Miami (Miami, FL), in the United States of America. Carbapenem resistance was confirmed by the Kirby Bauer disk diffusion method and the presence of the *bla*_KPC_ gene via polymerase chain reaction (PCR). Clonal complex 258 was confirmed by multiplex PCR. Multi-locus sequence typing of serial isolates was determined by *in silico* typing based on whole genome sequencing data. The minimum inhibitory concentrations of colistin were determined by the broth microdilution method as outlined by the Clinical and Laboratory Standards Institute(51).

Of the clinical KPC-Kp isolates collected, five genetically related serial KPC-Kp blood culture isolates - KLP0, KLP1, KLP2, KLP6, and KLP7 – were identified from a single critically-ill patient hospitalized at the University of Pittsburgh Medical Center and were utilized in subsequent experiments. The reference K2 serotype *Klebsiella pneumoniae* ATCC 43816 (American Type Culture Collection, Manassas, VA) was utilized in select experiments.

### Serum

Blood was obtained by routine venipuncture after informed consent (University of Pittsburgh STUDY19040346) from healthy volunteers. Serum donors were not taking any regular medications and abstained from aspirin and non-steroidal anti-inflammatory agents for 14 days prior to phlebotomy. Serum was separated in silica-coated plastic tubes (BD Vacutainer) then aliquoted and stored at -80°C. Serum was thawed at 37°C, pooled, and used immediately. Factor-depleted sera were obtained from Complement Technology (Tyler, Texas).

### Serum and polymyxin B resistance assay

Bacterial strains were grown overnight in tryptic soy broth (TSB) at 37°C and 250 RPM as previously described(18, 19). Briefly, overnight cultures were diluted 1:100 in TSB then re-incubated for 90-120 minutes to reach the early log phase of growth. Bacteria were pelleted by centrifugation at 2095 x *g* for 15 minutes, re-suspended in sterile culture-grade phosphate-buffered saline (PBS), and diluted to an OD_600_ = 0.2 (10^8^ colony forming units per milliliter) as per prior studies(19). Bacterial strains were then incubated at 37°C with 5% sterile TSB and 85% serum or 83 µM Polymyxin B (100 µg/mL in sterile PBS; Fresenius Kabi, Bad Homburg, Germany). Bacterial growth in serum was determined by measurement of OD_600_ every 60 minutes for 4 hours and quantified by the ratio of OD_600_ at 4 hours to OD_600_ at 0 hours. In select experiments, bacterial growth was confirmed by determining colony-forming units (CFU) via serial plating of bacteria.

### Whole genome sequencing

Genomic DNA from serial bloodstream isolates was extracted using a DNeasy Blood and Tissue Kit (Qiagen, Germantown, MD) from 1-mL bacterial cultures grown in Brain Heart Infusion (BHI) media as previously described(52). Next-generation sequencing libraries were prepared with a Nextera XT kit (Illumina, San Diego, CA), and libraries were sequenced on an Illumina MiSeq using 300-bp paired-end reads. Genomes were assembled with SPAdes v3.13.0, annotated with prokka version 1.14.5, and compared to one another with breseq version 0.33.2(53). The reference isolate KLP1 was also long-read sequenced on the Pacific Biosciences platform and used as a reference for identifying genetic variants in serial isolates from the same patient. Multi-locus sequence types and antimicrobial resistance genes were identified using online tools available through the Center for Genomic Epidemiology (http://www.genomicepidemiology.org/). The reference genome used, as well as Illumina read data for serial isolates newly sequenced in this study, will be submitted to NCBI under BioProject.

### Biofilm assay

Biofilm activity of clinical isolates was determined as previously described(54). Briefly, overnight cultures of bacterial strains were inoculated 1:100 in minimal M63 media supplemented with 0.5% casamino acids, 0.2% glucose, and 1 mM magnesium then aliquoted in triplicate into a clear, round-bottom 96 well plate and incubated in static conditions at 37°C for 24 hours. Bacterial growth was confirmed from a representative well on tryptic soy agar plates. Media was gently shaken out of the plate, which was then washed twice by gently submerging in de-ionized water and blotted dry on paper towels. Crystal violet (0.1%) was added in equal volume to all wells and incubated at room temperature for 15 minutes then gently washed by submersion in de-ionized water as above. The plate was inverted and dried for 4 hours at room temperature. Acetic acid (30%) was then added to each well for 15 minutes at room temperature to dissolve the crystal violet, which was transferred to a flat bottom 96-well plate for quantification by optical density at 550 nanometers.

### Osmotic tolerance assay

Serial KPC-Kp isolates were grown overnight in TSB, diluted 1:40 in TSB with 50% (volume/volume) sterile de-ionized water, and incubated at 37°C for 4 hours. Optical density at 600 nanometers was measured at 0 hours and 4 hours and osmotic tolerance was quantified by the ratio of OD_600_ at 4 hours to OD_600_ at 0 hours.

### Uronic acid quantification

Bacterial cultures were grown overnight, adjusted to similar concentrations, and plated on agar to confirm concentrations. Uronic acid content of the capsule was quantified based on published protocols with minor modifications(7, 55). Briefly, 0.5 mL of bacterial culture was mixed 1:1 with 1% Zwittergent 3-14 detergent in 100 mM citric acid (pH = 2.0) and then incubated at 50°C for 30 minutes with occasional mixing. The slurry was centrifuged at 16,000 x *g* for 5 minutes and 0.3 mL of supernatant was transferred to a new tube. To precipitate capsular polysaccharides, 1.2 mL of 100% culture grade ethanol was added, and the mixture was incubated at 4°C for 30 minutes. The suspension was centrifuged at 16,000 x *g* for 5 minutes.

The supernatant was discarded and the pellet was allowed to dry. The dry pellet was dissolved in 250 µL of 100 mM HCl overnight. The following morning, 20 µL of 4M ammonium sulfamate was mixed in 0.2 mL of the capsule extract on ice followed by the addition of 1 mL of 25 nM sodium tetraborate in sulfuric acid. The suspension was vortexed and placed at 100°C for 5 minutes then cooled to room temperature for 30 minutes. Thereafter, 40 µL of 0.15% m-hydroxybiphenyl was added and incubated for 15 minutes at room temperature followed immediately by measurement of optical density at 520 nanometers.

### Congo red agar plate assay

Brain heart infusion agar plates supplemented with 5% (weight/volume) sucrose and 0.08% (weight/volume) Congo Red were prepared as previously described(30). Bacteria from overnight cultures of serial KPC-Kp isolates were streaked onto each plate. Plates were left stationary at room temperature for 48 hours and then cataloged by digital photography.

### Transmission electron microscopy

Bacteria grown overnight on agar plates were stained per protocol(56). Briefly, colonies were fixed on agar with 2% paraformaldehyde and 2.5% glutaraldehyde in *0.1 M Na Cacodylate, 0.09 M sucrose, 0.01 M CaCl_2_.2H_2_O, 0.01 M MgCl .6H_2_O* (*Process Buffer*) supplemented with 1.55% L-lysine acetate (Acros) and 0.075% Ruthenium Red (Acros) for 20 minutes on ice. Cells were then incubated in fixative in Process Buffer with Ruthenium Red (without L-Lysine acetate) for 3 hours at 4°C. This was followed by 3 x 30-minute washes with Process Buffer with Ruthenium Red (no fixative) on ice. Cells were then post-fixed in 1% OsO_4_ with Ruthenium Red in Process Buffer for 1 hour on ice. Cells were washed 5 times for 10 minutes on ice in Process Buffer with Ruthenium Red. Colonies were removed from the agar plate and transferred to glass vials to complete processing for resin embedding.

For resin embedding, colonies were washed 3 times in PBS then dehydrated through a 30-100% ethanol series, then 100% propylene oxide, then infiltrated in 1:1 mixture of propylene oxide:Polybed 812 epoxy resin (Polysciences, Warrington, PA) for 1 hour. After several changes of 100% resin over 24 hours, colonies were embedded in molds and cured at 37°C overnight followed by additional hardening at 65°C for two more days. Semi-thin (300 nm) sections were heated onto glass slides, stained with 1% toluidine blue and imaged using light microscopy.

Ultrathin (60 nm) cross-sections of bacteria were collected on copper grids, stained with 1% uranyl acetate for 10 minutes, followed by 1% lead citrate for 7 minutes. Sections were viewed on a JEOL JEM 1400 FLASH transmission electron microscope (JEOL, Peabody MA) at 80 KV. Images were taken using a bottom mount AMT digital camera (Advanced Microscopy Techniques, Danvers, MA).

### Bacterial growth in complement C3-depleted sera

Select bacterial strains were grown to early log phase as described above and then incubated at 37°C with 5% sterile TSB and 85% serum or PBS control for 4 hours. In select experiments, bacteria were grown in sera depleted of complement C3 as well as in C3-depleted sera mixed 1:1 with healthy serum for 4 hours, as previously described(18). Bacterial growth was determined by quantifying CFU via serial plating of bacteria.

### Quantification of C3 and membrane attack complex binding by direct ELISA and flow cytometry

Serial bacterial isolates were grown to early log phase as described above then pelleted and re-suspended in cold sterile gelatin-veronal buffer with 5 mM magnesium chloride and EGTA (Complement Tech, Tyler, Texas) with 5% or 10% (volume/volume) normal healthy serum. Bacteria were incubated with serum at 37°C for 30 minutes with gentle mixing every 10 minutes. Cold sterile PBS was added to quench the reaction and the opsonized bacteria were pelleted at 2095 x *g* and washed x 3.

For the direct ELISA, opsonized bacteria were serially diluted and plated to confirm similar CFU counts. Bacteria (100 µL) were added in triplicate to a high-binding 96 well plate (Corning #9018) and incubated at 37°C without humidification to dry overnight. The bacteria were blocked with sterile-filtered 1% (weight/volume) bovine serum albumin for 2 hours and washed x 3 with 0.05% (volume/volume) Tween in PBS. To measure complement C3 binding, goat anti-human anti-C3 antibody (Catalog #A213, Complement Tech) was added at 1:200,000 dilution per manufacturer’s recommendation and incubated at room temperature for 1 hour, washed x 3, and then anti-goat horseradish peroxidase-conjugated antibody (Catalog #HAF017, R&D Systems, Minneapolis, MN) was added at 1:1000 dilution at room temperature for 1 hour. Membrane attack complex (MAC) binding was assessed by staining with rabbit anti-human C5b-C9 antibody (Catalog #204903, Sigma) at 1:1000 dilution at room temperature for 1 hour followed by washes x 3 then staining with goat anti-rabbit HRP-conjugated antibody (Catalog #204903, Sigma) at 1:5,000 dilution at room temperature for 1 hour. Following additional washes x 3, TMB substrate (Catalog #DY999, R&D Systems) was added for 10 minutes followed by equal parts stop solution per manufacturer’s instructions. The plate was measured for optical density at 450 nm and 540 nm. The OD_540_ was subtracted from the OD_450_ and the ratio of the corrected OD_450_ compared to the genetic reference isolate KLP1 was used to quantify binding.

For flow cytometry, after washing, opsonized bacteria were re-suspended in sterile PBS with 2% fetal bovine serum and stained with APC-conjugated anti-human C3b (Biolegend 846106) and FITC-conjugated anti-human C5b-C9 (LSBio C210266) for 30 minutes at 4°C. Cells were washed x 3, re-suspended in sterile PBS, and assayed on LSR-II flow cytometer (BD Biosciences, Franklin Lakes, NJ) at the University of Pittsburgh Unified Flow Core. Data were analyzed using FlowJo V10 (BD Biosciences).

### Immuno-electron microscopy

Bacteria were incubated with 5% or 10% normal human serum at 37°C for 30 minutes with gentle mixing every 10 minutes as described above. The reaction was quenched with cold sterile PBS and the opsonized bacteria were pelleted at 2095 x *g*, washed x 3, and then re-suspended in sterile PBS. Formvar-coated copper grids were placed coating side down onto puddles of treated bacterial suspensions for 5 minutes. Grids were then placed bacteria side down onto puddles of 2% paraformaldehyde in PBS for 10 minutes and then washed x 3 in PBS. Cells were blocked with 5% normal donkey serum in PBS for 30 min. Goat anti-C3 antibodies diluted 1:50 in 0.5% BSA, 0.1% glycine in PBS were added and incubated for 1 hour at room temperature, washed x 4 in PBS/BSA, and incubated with 6 nm colloidal gold conjugated donkey-anti-goat secondary antibodies (diluted 1:10 in PBS/BSA/glycine) for 1 hour. Cells were washed x 3 for 5 minutes with PBS/BSA/glycine then washed x 3 for 5 minutes with PBS and post-fixed in 2.5% glutaraldehyde in PBS for 10 minutes. Cells were washed x 3 with PBS then wicked dry with filter paper then grids were imaged on a JEOL JEM 1400 FLASH transmission electron microscope (JEOL, Peabody MA) at 80 kV. Images were taken using a bottom mount AMT digital camera (Advanced Microscopy Techniques, Danvers, MA).

### wcaJ allelic replacement

Approximately 500 base pairs (bps) of upstream and downstream sequences of the *wcaJ* gene were synthesized into the pUC57 vector from Genescript (Piscataway NJ). This vector was digested with BamHI and HindIII. The approximate 1000-bp fragment containing the flanking *wcaJ* sequences was gel extracted and cloned into the pMQ297 vector(57). Transformants were selected in Luria-Bertani (LB) broth plus hygromycin (140 µg/mL). To delete the chromosomal copy of *wcaJ* in the KLP1 isolate, the pMQ297 vector containing the upstream and downstream regions of *wcaJ* was introduced into the KLP1 isolate by electroporation. Transformants were selected for hygromycin resistance in LB broth plus hygromycin (140 µg/mL) grown at 42°C overnight. This allowed for integration of the plasmid into the bacterial genome. Merodiploids were temperature shifted to 30°C to promote removal of the plasmid from the chromosome. Individual colonies were tested for loss of hygromycin resistance, and hygromycin susceptible strains were assessed for deletion of *wcaJ* with PCR using forward (5’-GGTCTATGTCTTCGCTACTGC-3’) and reverse (5’-GCAGACAGCTCACGATTACG-3’) primers.

### wcaJ complementation

The capsule operon from KLP1 was cloned into the pBBR1-based cloning vector pMQ300(57) using yeast *in vivo* recombination(58) and placed under control of the *E. coli lac* promoter. The ∼21.3 kb *cps* operon from *galF* through *gnd* was amplified in two pieces of overlapping amplicons with primers that also directed recombination with pMQ300 using PrimeSTAR GXL high fidelity polymerase (Takara Bio). Primers to amplify the DNA were 5470 (5’-ttgtgagcggataacaatttcacacaggaaacagctGTGAAGATGAATATGGCGAATTTG-3’, lowercase letters direct recombination and uppercase priming) and 5429 (5’-gcatgatggaaggccctttcag-3’) for the *galF* containing amplicon and 5469 (5’-ccagtgccaagcttgcatgcctgcaggtcgactctagcaTTATTCCAGCCACTCGGTATG-3’) and 5474 (5’-GGTAATGATGCCAATTTGTTG-3’) for the *gnd* containing amplicon. The two amplicons and pMQ300 that had been linearized by SmaI (New England Biolabs) were used to transform *Saccharomyces cerevisiae* strain InvSc1 (Invitrogen). Uracil prototrophic transformants were pooled, plasmids were obtained and used to transform *E. coli* strain S17-1 lambda pir(59).

Candidates were tested by PCR, and the chosen plasmid was validated by whole plasmid sequencing (SNPsaurus, LLC), and dubbed pMQ787. The plasmid pMQ787 and vector control pMQ300 were moved into KLP1 and KLP1Δ*wcaJ* by conjugation with selection on LB agar with hygromycin (140 µg/mL) and kanamycin (50 µg/mL).

### Scanning electron microscopy

Twelve-millimeter round coverslips were treated with Cell-Tak (Corning). When the solution was dry, the treated side was placed on top of a bacterial colony for 10 min. Coverslips were removed, placed into individual wells of 24-well cluster plates and cells were fixed in 2.5% glutaraldehyde in PBS for 1 hour. Cells were washed 3 x 10 minutes in PBS then post-fixed for 1 hour in 1% aqueous OsO_4_. After 3 x 10 minutes PBS washes, cells were dehydrated in 30-100% ethanol series then 2 x 15 minutes in hexamethyldisilizane and then air dried. Coverslips were sputter-coated with 5 nm of gold/palladium alloy (Cressington) and viewed in a JEOL JSM-6330F scanning electron microscope (Peabody, MA) at 3 kV.

### Opsonized phagocytosis assay

RAW 264.7 macrophage cells (ATCC, Manassas, VA) were grown in sterile DMEM supplemented with 10% fetal bovine serum. After trypsin treatment, 0.25 x 10^6^ cells were placed in each well of a 24-well flat bottom tissue culture plate and grown overnight in fresh media at 37°C in humidified cell culture incubator with 5% carbon dioxide. The following morning, media was removed and replaced with fresh media with the addition of 10 µM cytochalasin D (Sigma, St. Louis, MO) to select wells with incubation for 1 hour prior to the addition of bacteria. Bacteria were grown into the early log phase as described above then pelleted and re-suspended in sterile PBS with 20% (volume/volume) normal healthy serum and placed on ice for 15 minutes to opsonize bacteria(60). Opsonized bacteria were pelleted at 2095 x *g*, re-suspended in cold sterile PBS, and added to each well at multiplicity of infection of 10:1. The tissue culture plate was centrifuged at 300 x *g* for 5 minutes to synchronize phagocytosis then incubated at 37°C for 1 hour. The medium was removed, and each well was washed with sterile PBS x 3 prior to incubation with 50 µM polymyxin B in DMEM at 37°C for 30 minutes to kill extracellular bacteria. Wells were washed with sterile PBS x 3 and cells were lysed with 0.25 mL of 1% (weight/volume) saponin (Sigma) for 10 minutes. Cell lysates were serially diluted and plated on tryptic soy agar overnight for CFU quantification.

### Mouse infection model

Wildtype (WT) C57BL/6J mice were purchased from Jackson Laboratory (Bar Harbor, Maine) and maintained using a protocol approved by the University of Pittsburgh Institutional Animal Care and Use Committee. Eight to sixteen-week old, age and sex-matched mice were utilized in all experiments. Alveolar macrophages were depleted with intra-tracheal inoculation of 0.5 mg room temperature clodronate (Encapsula #CLD-8901) in 0.1 mL volume or equivalent volume of empty liposome control 24 hours prior to infection. Mice were then intra-tracheally inoculated with 10^4^ CFU of strain KLP1 or KLP1Δ*wcaJ* in 0.1 mL. Necropsy was performed at 24 hours with collection of broncho-alveolar lavage fluid (BALF) and CFU counts as previously described (18, 61, 62). BALF cells were pelleted at 600 x *g* for 10 minutes at 4°C and the cell-free BALF supernatant was separated and stored at -80°C for subsequent assays. BALF cells were then processed for flow cytometry as described below.

### Flow cytometry of broncho-alveolar lavage

Flow cytometry on BALF cells was performed as described previously(63). Briefly, after pelleting and removal of cell-free BALF supernatant, the cells were incubated in ammonium-chloride-potassium (ACK) lysis buffer (150 mM NH4Cl, 10 mM KHCO3, 0.1 mM Na_2_EDTA in H2O, pH 7.2-7.4) for 5 minutes at room temperature to lyse red blood cells. Cells were then washed twice with PBS and stained with LIVE/DEAD Fixable Aqua Stain (Thermo Fisher) for 30 minutes at room temperature and protected from light. After this, cells were washed twice with PBS and resuspended in cell staining buffer (PBS with 2% newborn calf serum) containing the following anti-mouse antibodies (30 minutes at 4°C): CD45-Alexa Fluor 700 (clone: 30-F11, BD), CD11b-PE (clone M1/70, BD), Siglec-F-APC-Cy7 (clone: E50-2440, BD), CD11c-PE-Cy7 (clone HL3, BD), CD64-BV650 (clone: X54-5/7.1, BD), Ly6G-APC (clone 1A8, BD). Flow cytometry analysis was performed using an LSR Fortessa Cell Analyzer (BD) at the University of Pittsburgh Unified Flow Core. Absolute cell counts (cells/mL) were determined using CountBright Absolute Counting Beads (Thermo Fisher). Data were analyzed using FlowJo version 10.8.0 (BD).

### Ex vivo broncho-alveolar lavage fluid opsonization assay and protein content

We modified an existing immunoassay for complement component deposition to measure complement C3 deposition by BALF on bacteria(64). The KLP1Δ*wcaJ* strain was grown to log phase and 10^8^ CFU/mL bacteria were plated at 0.1 mL volume in each well of a 96-well high binding plate (Corning #9018). The plates were dried uncovered overnight at 37°C without humidification. The following day, bacteria were washed x 3, blocked with sterile PBS with 1% BSA for a minimum of 1 hour, and then washed x 3. The wells were incubated at 37°C and agitated at 50 RPM for 1 hour with either 0.1 mL of sterile PBS, 20% vol/vol mouse serum from uninfected mice with or without 5 mM EDTA to block complement function, or BALF from uninfected mice or BALF collected at necropsy from the KLP1Δ*wcaJ* lung infection experiment described above. Following incubation, complement activity was quenched with the addition of 0.2 mL of ice-cold sterile PBS followed by washing x 3. Wells were coated with 0.1 mL of goat anti-mouse C3 (Complement Tech #M213, 1:500 dilution per manufacturer recommendations) for 1 hour at room temperature followed by washing x 5. Anti-goat horseradish peroxidase-conjugated antibody (Catalog #HAF017, R&D Systems, Minneapolis, MN) at 1:1000 dilution was added and incubated at room temperature for 1 hour followed by washing x 5. TMB substrate (Catalog #DY999, R&D Systems) was added for 10 minutes followed by stop solution per manufacturer’s instructions. The optical density at 450 nm and 540 nm was measured. The OD_540_ was subtracted from the OD_450_ followed by subtraction of background calculated as the median of values from PBS-coated wells. BALF protein content was quantified by BCA assay as previously described(62).

### Human whole blood opsono-phagocytosis assay

The protocol for whole blood opsono-phagocytosis was adapted from prior publications (33, 34). Bacterial isolates were grown to log-phase then diluted to 10^8^ CFU/mL and heat-killed at 60°C for 1 hour then washed once and re-suspended in 100 mM sodium bicarbonate. The heat-killed isolates were labeled with 1mM pHrodo green dye per manufacturer instructions (Invitrogen # P36013). Blood was withdrawn from a single healthy donor without prior medical history or medications for 14 days using a 21-gauge needle and immediately added to a tube with sterile lepiriudin (Celgene) to final concentration of 50 µg/mL and gently rocked for 10 seconds. Thereafter, the lepirudin-treated blood was immediately divided and sterile compstatin (Tocris #2585), or an equivalent volume of sterile PBS vehicle, was added to final concentration of 167 µM. The blood was then further sub-divided and 10^7^ CFU of each heat-killed, pHrodo-labeled isolate was added to an individual tube. Immediately, the tubes were placed into a pre-warmed incubator at 37°C and rotated for 30 minutes. The reaction was immediately quenched by the addition of 10 mM of ice-cold sterile EDTA followed by centrifugation at 300 g for 10 minutes at 4°C. The blood cell pellet was then re-suspended in ACK lysis buffer for 5 minutes at room temperature to lyse red blood cells then centrifuged at 300 g for 10 minutes and washed in PBS twice. The cell pellet was resuspended in PBS with 1% BSA and immediately analyzed on BD FACScalibur using forward and side-scatter gating.

### Statistical analysis

Two-tailed Mann-Whitney U test was used when comparing 2 groups, and a Kruskal-Wallis test with Dunn’s post-hoc test for multiple comparisons were used when comparing >2 groups, unless otherwise indicated. P<0.05 was considered significant. All statistics were performed using GraphPad Prism V9 (La Jolla, CA).

---

**Supplemental Figure 1.**
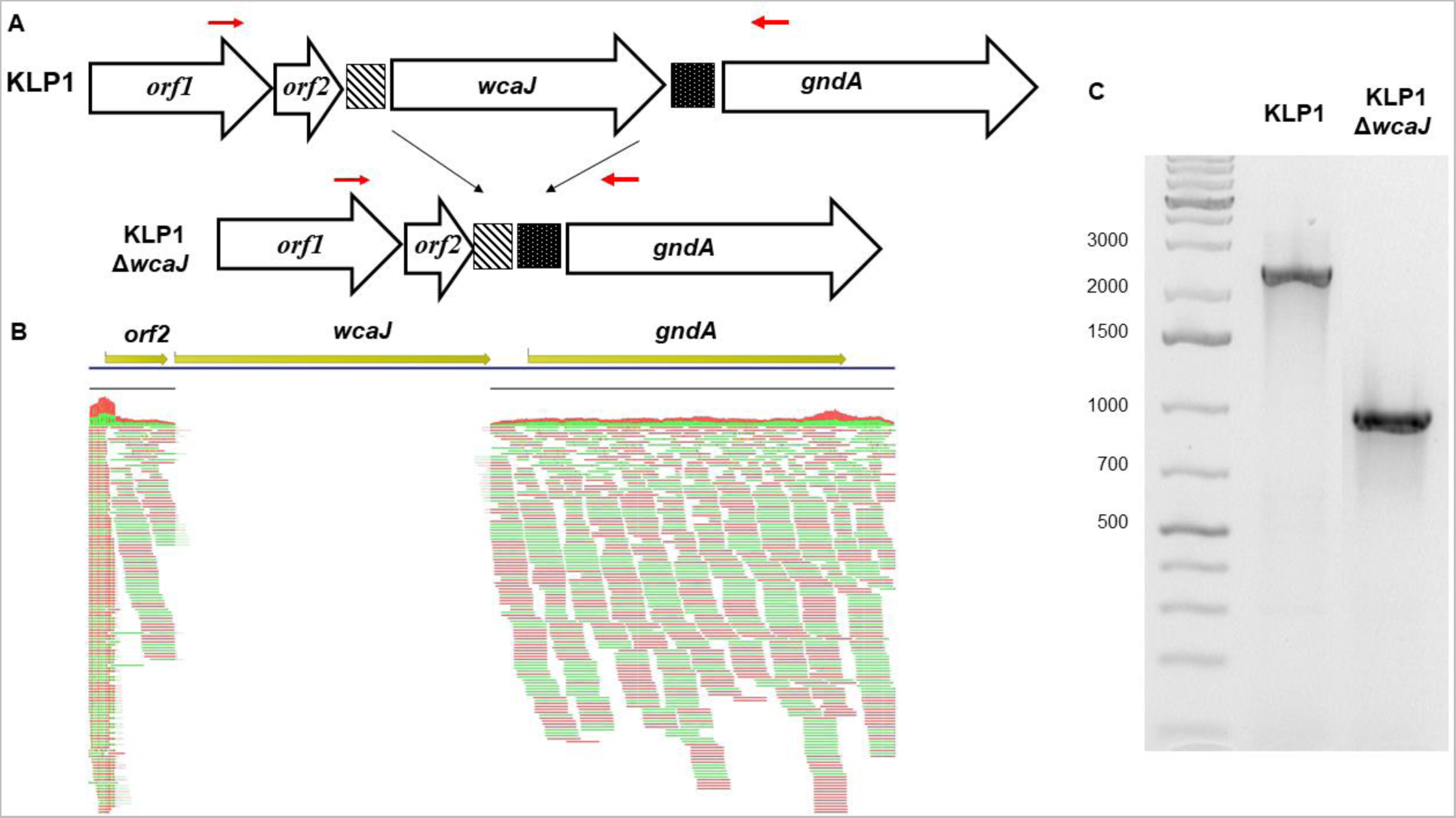
Deletion of wcaJ in isogenic reference isolate. A) Schematic demonstrating *wcaJ* genetic environment in KLP1 and KLP1*ΔwcaJ* isolates. The direction of transcription of genes is shown by white arrows. Striped and black squares show intergenic sequences. Red arrows show the approximate location of PCR primers used to confirm *wcaJ* deletion. (B) Image from CLC showing absence of the Illumina read coverage of *wcaJ* gene in KLP1Δ*wcaJ* isolate, consistent with gene deletion. (C) PCR image confirming the presence of *wcaJ* in KLP1 isolates, and deletion of *wcaJ* in KLP1Δ*wcaJ*. Abbreviations: orf= open reading frame; gndA= NADP-dependent phosphogluconate dehydrogenase

**Supplemental Figure 2.**
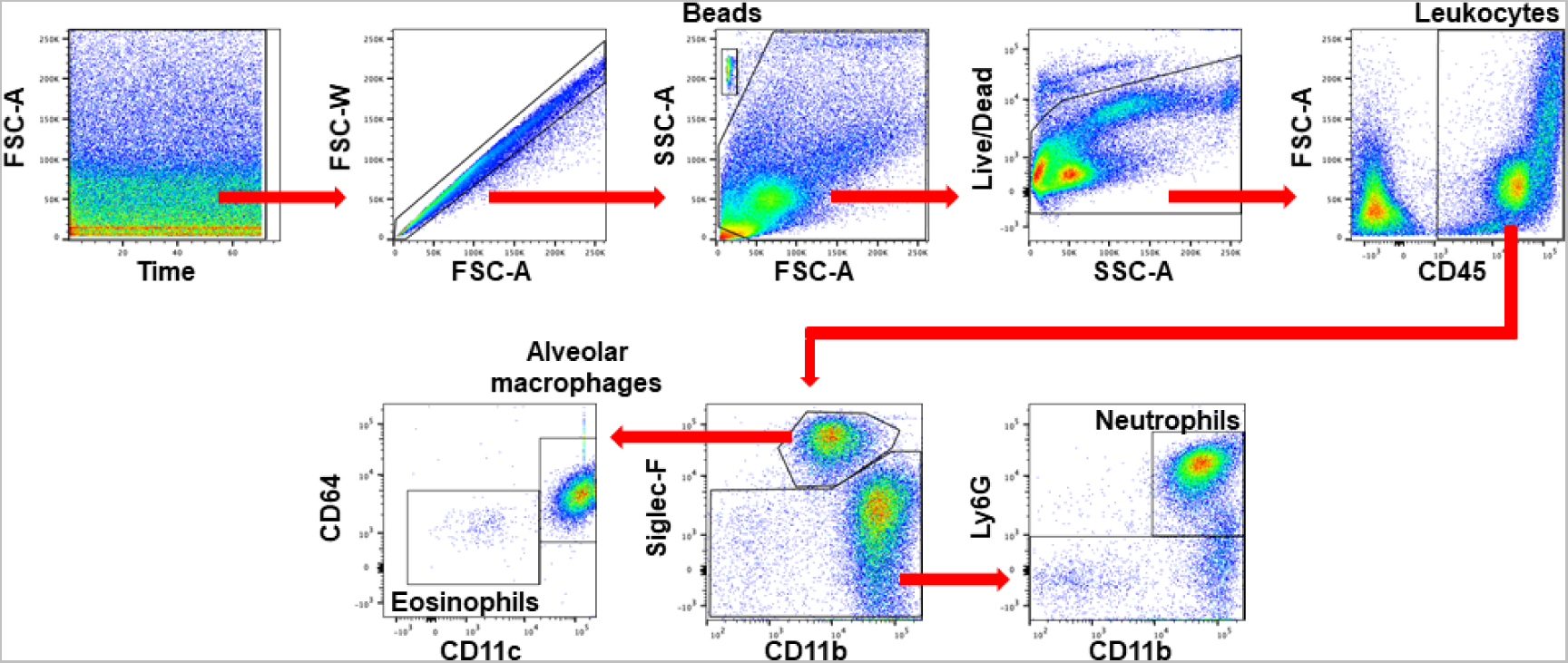
**Flow cytometry gating strategy for broncho-alveolar lavage fluid cells.**

## Bibliography

1. Gonzalez-Ferrer S, Peñaloza HF, Budnick JA, Bain WG, Nordstrom HR, Lee JS, Van Tyne D. 2021. Finding Order in the Chaos: Outstanding Questions in Klebsiella pneumoniae Pathogenesis. Infect Immun 89.

2. Gu D, Dong N, Zheng Z, Lin D, Huang M, Wang L, Chan EW-C, Shu L, Yu J, Zhang R, Chen S. 2018. A fatal outbreak of ST11 carbapenem-resistant hypervirulent Klebsiella pneumoniae in a Chinese hospital: a molecular epidemiological study. Lancet Infect Dis 18:37–46.

3. Munoz-Price LS, Poirel L, Bonomo RA, Schwaber MJ, Daikos GL, Cormican M, Cornaglia G, Garau J, Gniadkowski M, Hayden MK, Kumarasamy K, Livermore DM, Maya JJ, Nordmann P, Patel JB, Paterson DL, Pitout J, Villegas MV, Wang H, Woodford N, Quinn JP. 2013. Clinical epidemiology of the global expansion of Klebsiella pneumoniae carbapenemases. Lancet Infect Dis 13:785–796.

4. Borer A, Saidel-Odes L, Riesenberg K, Eskira S, Peled N, Nativ R, Schlaeffer F, Sherf M. 2009. Attributable mortality rate for carbapenem-resistant Klebsiella pneumoniae bacteremia. Infect Control Hosp Epidemiol 30:972–976.

5. Ben-David D, Kordevani R, Keller N, Tal I, Marzel A, Gal-Mor O, Maor Y, Rahav G. 2012. Outcome of carbapenem resistant Klebsiella pneumoniae bloodstream infections. Clin Microbiol Infect 18:54–60.

6. Fraenkel-Wandel Y, Raveh-Brawer D, Wiener-Well Y, Yinnon AM, Assous MV. 2016. Mortality due to blaKPC Klebsiella pneumoniae bacteraemia. J Antimicrob Chemother 71:1083–1087.

7. Ernst CM, Braxton JR, Rodriguez-Osorio CA, Zagieboylo AP, Li L, Pironti A, Manson AL, Nair AV, Benson M, Cummins K, Clatworthy AE, Earl AM, Cosimi LA, Hung DT. 2020. Adaptive evolution of virulence and persistence in carbapenem-resistant Klebsiella pneumoniae. Nat Med 26:705–711.

8. DeLeo FR, Kobayashi SD, Porter AR, Freedman B, Dorward DW, Chen L, Kreiswirth BN. 2017. Survival of Carbapenem-Resistant Klebsiella pneumoniae Sequence Type 258 in Human Blood. Antimicrob Agents Chemother 61.

9. Tzouvelekis LS, Miriagou V, Kotsakis SD, Spyridopoulou K, Athanasiou E, Karagouni E, Tzelepi E, Daikos GL. 2013. KPC-producing, multidrug-resistant Klebsiella pneumoniae sequence type 258 as a typical opportunistic pathogen. Antimicrob Agents Chemother 57:5144–5146.

10. Ahn D, Peñaloza H, Wang Z, Wickersham M, Parker D, Patel P, Koller A, Chen EI, Bueno SM, Uhlemann A-C, Prince A. 2016. Acquired resistance to innate immune clearance promotes Klebsiella pneumoniae ST258 pulmonary infection. JCI Insight 1:e89704.

11. Peñaloza HF, Noguera LP, Ahn D, Vallejos OP, Castellanos RM, Vazquez Y, Salazar-Echegarai FJ, González L, Suazo I, Pardo-Roa C, Salazar GA, Prince A, Bueno SM. 2019. Interleukin-10 Produced by Myeloid-Derived Suppressor Cells Provides Protection to Carbapenem-Resistant Klebsiella pneumoniae Sequence Type 258 by Enhancing Its Clearance in the Airways. Infect Immun 87.

12. Siu LK, Yeh K-M, Lin J-C, Fung C-P, Chang F-Y. 2012. *Klebsiella pneumoniae* liver abscess: a new invasive syndrome. Lancet Infect Dis 12:881–887.

13. Harada S, Doi Y. 2018. Hypervirulent *Klebsiella pneumoniae*: a Call for Consensus Definition and International Collaboration. J Clin Microbiol 56.

14. Liu YC, Cheng DL, Lin CL. 1986. *Klebsiella pneumoniae* liver abscess associated with septic endophthalmitis. Arch Intern Med 146:1913–1916.

15. Casadevall A, Pirofski L. 2001. Host-pathogen interactions: the attributes of virulence. J Infect Dis 184:337–344.

16. Domenico P, Salo RJ, Cross AS, Cunha BA. 1994. Polysaccharide capsule-mediated resistance to opsonophagocytosis in Klebsiella pneumoniae. Infect Immun 62:4495–4499.

17. March C, Cano V, Moranta D, Llobet E, Pérez-Gutiérrez C, Tomás JM, Suárez T, Garmendia J, Bengoechea JA. 2013. Role of bacterial surface structures on the interaction of Klebsiella pneumoniae with phagocytes. PLoS ONE 8:e56847.

18. Bain W, Li H, van der Geest R, Moore SR, Olonisakin TF, Ahn B, Papke E, Moghbeli K, DeSensi R, Rapport S, Saul M, Hulver M, Xiong Z, Mallampalli RK, Ray P, Morris A, Ma L, Doi Y, Zhang Y, Kitsios GD, Kulkarni HS, McVerry BJ, Ferreira VP, Nouraie M, Lee JS. 2020. Increased Alternative Complement Pathway Function and Improved Survival during Critical Illness. Am J Respir Crit Care Med 202:230–240.

19. Olonisakin TF, Li H, Xiong Z, Kochman EJK, Yu M, Qu Y, Hulver M, Kolls JK, St Croix C, Doi Y, Nguyen M-H, Shanks RMQ, Mallampalli RK, Kagan VE, Ray A, Silverstein RL, Ray P, Lee JS. 2016. CD36 Provides Host Protection Against Klebsiella pneumoniae Intrapulmonary Infection by Enhancing Lipopolysaccharide Responsiveness and Macrophage Phagocytosis. J Infect Dis 214:1865–1875.

20. Broug-Holub E, Toews GB, van Iwaarden JF, Strieter RM, Kunkel SL, Paine R, Standiford TJ. 1997. Alveolar macrophages are required for protective pulmonary defenses in murine Klebsiella pneumonia: elimination of alveolar macrophages increases neutrophil recruitment but decreases bacterial clearance and survival. Infect Immun 65:1139–1146.

21. Loraine J, Heinz E, De Sousa Almeida J, Milevskyy O, Voravuthikunchai SP, Srimanote P, Kiratisin P, Thomson NR, Taylor PW. 2018. Complement Susceptibility in Relation to Genome Sequence of Recent Klebsiella pneumoniae Isolates from Thai Hospitals. mSphere 3.

22. Kobayashi SD, Porter AR, Freedman B, Pandey R, Chen L, Kreiswirth BN, DeLeo FR. 2018. Antibody-Mediated Killing of Carbapenem-Resistant ST258 Klebsiella pneumoniae by Human Neutrophils. MBio 9.

23. Casalino M, Prosseda G, Barbagallo M, Iacobino A, Ceccarini P, Latella MC, Nicoletti M, Colonna B. 2010. Interference of the CadC regulator in the arginine-dependent acid resistance system of Shigella and enteroinvasive E. coli. Int J Med Microbiol 300:289–295.

24. Yaron A. 1976. Dipeptidyl carboxypeptidase from Escherichia coli. Meth Enzymol 45:599–610.

25. Poirel L, Jayol A, Bontron S, Villegas M-V, Ozdamar M, Türkoglu S, Nordmann P. 2015. The mgrB gene as a key target for acquired resistance to colistin in Klebsiella pneumoniae. J Antimicrob Chemother 70:75–80.

26. Kato A, Tanabe H, Utsumi R. 1999. Molecular characterization of the PhoP-PhoQ two-component system in Escherichia coli K-12: identification of extracellular Mg2+-responsive promoters. J Bacteriol 181:5516–5520.

27. Cannatelli A, Giani T, D’Andrea MM, Di Pilato V, Arena F, Conte V, Tryfinopoulou K, Vatopoulos A, Rossolini GM, COLGRIT Study Group. 2014. MgrB inactivation is a common mechanism of colistin resistance in KPC-producing Klebsiella pneumoniae of clinical origin. Antimicrob Agents Chemother 58:5696–5703.

28. Kidd TJ, Mills G, Sá-Pessoa J, Dumigan A, Frank CG, Insua JL, Ingram R, Hobley L, Bengoechea JA. 2017. A *Klebsiella pneumoniae* antibiotic resistance mechanism that subdues host defences and promotes virulence. EMBO Mol Med 9:430–447.

29. Short FL, Di Sario G, Reichmann NT, Kleanthous C, Parkhill J, Taylor PW. 2020. Genomic Profiling Reveals Distinct Routes To Complement Resistance in Klebsiella pneumoniae. Infect Immun 88.

30. Van Tyne D, Ciolino JB, Wang J, Durand ML, Gilmore MS. 2016. Novel Phagocytosis-Resistant Extended-Spectrum β-Lactamase-Producing Escherichia coli From Keratitis. JAMA Ophthalmol 134:1306–1309.

31. Doorduijn DJ, Rooijakkers SHM, van Schaik W, Bardoel BW. 2016. Complement resistance mechanisms of Klebsiella pneumoniae. Immunobiology 221:1102–1109.

32. de Astorza B, Cortés G, Crespí C, Saus C, Rojo JM, Albertí S. 2004. C3 promotes clearance of Klebsiella pneumoniae by A549 epithelial cells. Infect Immun 72:1767–1774.

33. Mollnes TE, Brekke O-L, Fung M, Fure H, Christiansen D, Bergseth G, Videm V, Lappegård KT, Köhl J, Lambris JD. 2002. Essential role of the C5a receptor in E coli-induced oxidative burst and phagocytosis revealed by a novel lepirudin-based human whole blood model of inflammation. Blood 100:1869–1877.

34. Bexborn F, Engberg AE, Sandholm K, Mollnes TE, Hong J, Nilsson Ekdahl K. 2009. Hirudin versus heparin for use in whole blood in vitro biocompatibility models. J Biomed Mater Res A 89:951–959.

35. Deleo FR, Chen L, Porcella SF, Martens CA, Kobayashi SD, Porter AR, Chavda KD, Jacobs MR, Mathema B, Olsen RJ, Bonomo RA, Musser JM, Kreiswirth BN. 2014. Molecular dissection of the evolution of carbapenem-resistant multilocus sequence type 258 Klebsiella pneumoniae. Proc Natl Acad Sci USA 111:4988–4993.

36. Tan YH, Chen Y, Chu WHW, Sham L-T, Gan Y-H. 2020. Cell envelope defects of different capsule-null mutants in K1 hypervirulent Klebsiella pneumoniae can affect bacterial pathogenesis. Mol Microbiol 113:889–905.

37. Rendueles O. 2020. Deciphering the role of the capsule of Klebsiella pneumoniae during pathogenesis: A cautionary tale. Mol Microbiol 113:883–888.

38. Stevenson G, Andrianopoulos K, Hobbs M, Reeves PR. 1996. Organization of the Escherichia coli K-12 gene cluster responsible for production of the extracellular polysaccharide colanic acid. J Bacteriol 178:4885–4893.

39. Pal S, Verma J, Mallick S, Rastogi SK, Kumar A, Ghosh AS. 2019. Absence of the glycosyltransferase WcaJ in Klebsiella pneumoniae ATCC13883 affects biofilm formation, increases polymyxin resistance and reduces murine macrophage activation. Microbiology (Reading, Engl) 165:891–904.

40. Jousset AB, Bonnin RA, Rosinski-Chupin I, Girlich D, Cuzon G, Cabanel N, Frech H, Farfour E, Dortet L, Glaser P, Naas T. 2018. A 4.5-Year Within-Patient Evolution of a Colistin-Resistant Klebsiella pneumoniae Carbapenemase-Producing K. pneumoniae Sequence Type 258. Clin Infect Dis 67:1388–1394.

41. Merino S, Camprubí S, Albertí S, Benedí VJ, Tomás JM. 1992. Mechanisms of Klebsiella pneumoniae resistance to complement-mediated killing. Infect Immun 60:2529–2535.

42. Cortés G, Borrell N, de Astorza B, Gómez C, Sauleda J, Albertí S. 2002. Molecular analysis of the contribution of the capsular polysaccharide and the lipopolysaccharide O side chain to the virulence of Klebsiella pneumoniae in a murine model of pneumonia. Infect Immun 70:2583– 2590.

43. Formosa C, Herold M, Vidaillac C, Duval RE, Dague E. 2015. Unravelling of a mechanism of resistance to colistin in Klebsiella pneumoniae using atomic force microscopy. J Antimicrob Chemother 70:2261–2270.

44. Jensen TS, Opstrup KV, Christiansen G, Rasmussen PV, Thomsen ME, Justesen DL, Schønheyder HC, Lausen M, Birkelund S. 2020. Complement mediated Klebsiella pneumoniae capsule changes. Microbes Infect 22:19–30.

45. Doorduijn DJ, Bardoel BW, Heesterbeek DAC, Ruyken M, Benn G, Parsons ES, Hoogenboom BW, Rooijakkers SHM. 2020. Bacterial killing by complement requires direct anchoring of membrane attack complex precursor C5b-7. PLoS Pathog 16:e1008606.

46. Heesterbeek DA, Bardoel BW, Parsons ES, Bennett I, Ruyken M, Doorduijn DJ, Gorham RD, Berends ET, Pyne AL, Hoogenboom BW, Rooijakkers SH. 2019. Bacterial killing by complement requires membrane attack complex formation via surface-bound C5 convertases. EMBO J 38.

47. Dudkina NV, Spicer BA, Reboul CF, Conroy PJ, Lukoyanova N, Elmlund H, Law RHP, Ekkel SM, Kondos SC, Goode RJA, Ramm G, Whisstock JC, Saibil HR, Dunstone MA. 2016. Structure of the poly-C9 component of the complement membrane attack complex. Nat Commun 7:10588.

48. Joiner KA. 1988. Complement evasion by bacteria and parasites. Annu Rev Microbiol 42:201– 230.

49. Mueller-Ortiz SL, Drouin SM, Wetsel RA. 2004. The alternative activation pathway and complement component C3 are critical for a protective immune response against Pseudomonas aeruginosa in a murine model of pneumonia. Infect Immun 72:2899–2906.

50. Ben-David I, Price SE, Bortz DM, Greineder CF, Cohen SE, Bauer AL, Jackson TL, Younger JG. 2005. Dynamics of intrapulmonary bacterial growth in a murine model of repeated microaspiration. Am J Respir Cell Mol Biol 33:476–482.

51. Clinical and Laboratory Standards Institute. 2018. Methods for Dilution Antimicrobial Susceptibility Tests for Bacteria That Grow Aerobically.

52. Chilambi GS, Nordstrom HR, Evans DR, Kowalski RP, Dhaliwal DK, Jhanji V, Shanks RMQ, Van Tyne D. 2021. Genomic and phenotypic diversity of Enterococcus faecalis isolated from endophthalmitis. PLoS ONE 16:e0250084.

53. Deatherage DE, Barrick JE. 2014. Identification of mutations in laboratory-evolved microbes from next-generation sequencing data using breseq. Methods Mol Biol 1151:165–188.

54. Zupetic J, Peñaloza HF, Bain W, Hulver M, Mettus R, Jorth P, Doi Y, Bomberger J, Pilewski J, Nouraie M, Lee JS. 2021. Elastase Activity From Pseudomonas aeruginosa Respiratory Isolates and ICU Mortality. Chest 160:1624–1633.

55. Domenico P, Schwartz S, Cunha BA. 1989. Reduction of capsular polysaccharide production in Klebsiella pneumoniae by sodium salicylate. Infect Immun 57:3778–3782.

56. Birkhead M, Ganesh K, Ndlangisa K, Koornhof H. 2017. Transmission electron microscopy protocols for capsule visualisation in pathogenic respiratory and meningeal bacteria, p. 628–39. In Microscopy and imaging science : practical approaches to applied research and education.

57. Kalivoda EJ, Horzempa J, Stella NA, Sadaf A, Kowalski RP, Nau GJ, Shanks RMQ. 2011. New vector tools with a hygromycin resistance marker for use with opportunistic pathogens. Mol Biotechnol 48:7–14.

58. Shanks RMQ, Caiazza NC, Hinsa SM, Toutain CM, O’Toole GA. 2006. *Saccharomyces cerevisiae*-based molecular tool kit for manipulation of genes from gram-negative bacteria. Appl Environ Microbiol 72:5027–5036.

59. Miller VL, Mekalanos JJ. 1988. A novel suicide vector and its use in construction of insertion mutations: osmoregulation of outer membrane proteins and virulence determinants in Vibrio cholerae requires toxR. J Bacteriol 170:2575–2583.

60. Zhao Y, Olonisakin TF, Xiong Z, Hulver M, Sayeed S, Yu MT, Gregory AD, Kochman EJ, Chen BB, Mallampalli RK, Sun M, Silverstein RL, Stolz DB, Shapiro SD, Ray A, Ray P, Lee JS. 2015. Thrombospondin-1 restrains neutrophil granule serine protease function and regulates the innate immune response during Klebsiella pneumoniae infection. Mucosal Immunol 8:896–905.

61. Qu Y, Olonisakin T, Bain W, Zupetic J, Brown R, Hulver M, Xiong Z, Tejero J, Shanks RM, Bomberger JM, Cooper VS, Zegans ME, Ryu H, Han J, Pilewski J, Ray A, Cheng Z, Ray P, Lee JS. 2018. Thrombospondin-1 protects against pathogen-induced lung injury by limiting extracellular matrix proteolysis. JCI Insight 3.

62. Bain W, Olonisakin T, Yu M, Qu Y, Hulver M, Xiong Z, Li H, Pilewski J, Mallampalli RK, Nouraie M, Ray A, Ray P, Cheng Z, Shanks RMQ, St Croix C, Silverstein RL, Lee JS. 2019. Platelets inhibit apoptotic lung epithelial cell death and protect mice against infection-induced lung injury. Blood Adv 3:432–445.

63. Peñaloza HF, Olonisakin TF, Bain WG, Qu Y, van der Geest R, Zupetic J, Hulver M, Xiong Z, Newstead MW, Zou C, Alder JK, Ybe JA, Standiford TJ, Lee JS. 2021. Thrombospondin-1 Restricts Interleukin-36γ-Mediated Neutrophilic Inflammation during Pseudomonas aeruginosa Pulmonary Infection. MBio 12.

64. Ben Nasr A, Klimpel GR. 2008. Subversion of complement activation at the bacterial surface promotes serum resistance and opsonophagocytosis of Francisella tularensis. J Leukoc Biol 84:77– 85.

